# On deleterious mutations in perennials: inbreeding depression, mutation load and life-history evolution

**DOI:** 10.1101/865220

**Authors:** Thomas Lesaffre, Sylvain Billiard

## Abstract

In Angiosperms, perennials typically present much higher levels of inbreeding depression than annuals. One hypothesis to explain this pattern stems from the observation that inbreeding depression is expressed across multiple life stages in Angiosperms. It posits that increased inbreeding depression in more long-lived species could be explained by differences in the way mutations affect fitness in these species, through the life stages at which they are expressed. In this study, we investigate this hypothesis. We combine a physiological growth model and multilocus population genetics approaches to describe a full genotype-to-phenotype-to-fitness map. We study the behaviour of mutations affecting growth or survival, and explore their consequences in terms of inbreeding depression and mutation load. Although our results only agree with empirical data within a narrow range of conditions, we argue that they may point us towards the type of traits susceptible to underlie inbreeding depression in long-lived species, that is traits under sufficiently strong selection, on which selection decreases sharply as life expectancy increases. Then, we study the role deleterious mutations maintained at mutation-selection balance may play in the coevolution between growth and survival strategies.

**Description of the manuscript:** The main text of the manuscript, excluding captions and headers, is 5712 words long. There are 4 figures in the main text, numbered from 1 to 4. In the present file, pages 1 to 33 correspond to the main text (including title page, abstract and litterature cited), while the remaining pages (33 to 72) correspond to appendices. There are 5 sections in Appendices, which are all available at the end of the manuscript file. There are 12 figures in Appendices, numbered from S1 to S12.

## 1 Introduction

Perennials, which make up the majority of Angiosperms (~ 70%, Munoz et al., 2016), typically present much higher levels of inbreeding depression than annuals. Indeed, meta-analyses found inbreeding depression to span from *δ* ≈ 0.2 on average in short-lived herbaceous species to *δ* ≈ 0.5 in long-lived herbaceous species and shrubs, and *δ* ≈ 0.6 in woody species (Duminil et al., 2009; Angeloni et al., 2011). Inbreeding depression, defined as the reduction in fitness of inbred relative to outbred individuals, is thought to be mainly due to the increased homozygosity of inbred individuals for recessive deleterious mutations segregating at low frequencies in populations (Charlesworth and Charlesworth, 1987; Charlesworth and Willis, 2009). Why inbreeding depression is higher in more long-lived species is still poorly understood, despite the potential significance of this pattern for important evolutionary questions, such as mating systems or dispersal rates evolution (Barrett and Harder, 1996; Roze and Rousset, 2005; Epinat and Lenormand, 2009; Duputié and Massol, 2013), and for more applied issues, as many cultivated species are perennial (e.g. fruit trees in general) and efforts are being made to develop perennial grain crops (DeHaan and Van Tassel, 2014).

Two hypotheses have been proposed to explain this pattern. The first hypothesis was formally put forward by Scofield and Schultz (2006). In plants, mutations occurring in somatic tissues may be passed onto the offspring, because they do not have a segregated germline. Thus, Scofield and Schultz (2006) proposed that more long-lived species may accumulate more somatic mutations as they grow and transmit them to their offspring, thereby generating the increase in inbreeding depression observed in such species. Phenotypic data in a long-lived clonal shrub (*Vaccinium angustifolium*, Bobiwash et al., 2013), and genomic results in *Quercus robur* (Plomion et al., 2018) demonstrated that some somatic mutations can indeed be passed onto the offspring. However, the number of detected heritable somatic mutations is low (Schmid-Siegert et al., 2017; Plomion et al., 2018), and recent studies have concluded that the number of cell divisions from embryonic cells to gametes production may be much lower than previously thought (Burian et al., 2016; Watson et al., 2016; Burian et al., 2016; Schmid-Siegert et al., 2017; Lanfear, 2018), due for instance to early specification and quiescence mechanisms of axillary meristems cells, resulting in little opportunity for heritable mutations to accumulate during plant growth. Furthermore, intraorganismal selection is expected to efficiently purge deleterious somatic mutations, resulting in little to no somatically generated mutation load at the population level (Otto and Orive, 1995). Hence, although somatic mutations can be inherited and contribute to the mutation load in plants, their relative significance compared with meiotic mutations remains unclear (Schoen and Schultz, 2019).

The second hypothesis stems from the observation that inbreeding depression is typically expressed across multiple life stages in Angiosperms (Husband and Schemske, 1996; Winn et al., 2011; Angeloni et al., 2011). It posits that increased inbreeding depression in more long-lived species could be explained by the fact that mutations, regardless of whether they are produced during mitosis or meiosis, may differ in the way they affect fitness in annual and perennial populations, through the life stages at which they are expressed. Most theoretical studies of the mutation load focused on the case of mutations affecting fitness on a strictly linear fitness landscape (that is, fitness is the trait, e.g. Charlesworth et al., 1990; Roze, 2015), or through an abstract trait (or set of traits) under stabilising selection on a gaussian fitness landscape (e.g. Roze and Blanckaert, 2014; Abu Awad and Roze, 2018). In these cases, inbreeding depression in annual and perennial populations is not expected to differ (Charlesworth, 1980). On the other hand, the dynamics of mutations affecting other aspects of individuals’ life cycle, such as survival or growth, were seldom investigated. Morgan (2001) investigated the dynamics of mutations affecting survival between mating events in a perennial populations. They concluded that inbreeding depression should sharply decrease as life expectancy increases, and even become negative for long-lived species (outbreeding depression). However, Morgan (2001) studied mutations with a strong effect on fitness, and assumed no age-structure, that is, individuals did not differ in fecundity or survival probability with age. Such variations with respect to age in both survival and fecundity are yet observed in perennials. Indeed, while juveniles typically suffer from very high mortality rates, established individuals tend to experience rather low mortalities with particularly slow senescence (Petit and Hampe, 2006). Furthermore, fecundity usually increases dramatically with age in perennials (Franco and Silvertown, 1996), due to the positive scaling of reproductive output with size in plants (Klinkhamer et al., 1985; Weiner et al., 2009). Mutations slowing their bearer’s growth could therefore play a role in generating higher inbreeding depression in more long-lived species, as growth delays impact individuals’ fecundities negatively and with varying intensity with respect to age or size. This latter aspect of age-structuration in perennials was, to our knowledge, never tackled theoretically.

The present study aims to study the second hypothesis, that is, we investigate the behaviour of meiotic mutations affecting fitness differently with respect to life-history, putting aside somatic mutations accumulation. Namely, we study meiotic mutations affecting growth or survival in a partially selfing population, in which individuals grow as they age and fecundity is proportional to size, but survival between mating events is assumed to not depend on age. We combine a physiological growth model (West et al., 2001) and multilocus population genetics approaches (Barton and Turelli, 1991; Kirkpatrick et al., 2002) in order to describe a full genotype-to-phenotype-to-fitness map, where the fitness landscape emerges from developmental processes instead of being assumed *a priori*. We study the behaviour of different types of mutations affecting growth or survival, and explore their consequences in terms of inbreeding depression and mutation load (Crow, 1958). Then, we show the role deleterious mutations maintained at mutation-selection balance may play in the coevolution between growth and survival strategies.

## 2 Model outline and methods

### Demography

We consider a large population of diploid hermaphrodites, which may survive from one mating season to another with a probability *S*, assumed to be constant with respect to age and size. If they survive, individuals grow between mating events, following a physiological growth model described briefly in the next paragraph. If they die, juveniles, which are assumed to be produced in large excess compared with the resources available for establishment, are recruited to replace the dead, so that population size is kept constant (Fig. 1). We assume that juveniles are produced each year and do not survive to the next mating season if they are not recruited (no dormancy). Each juvenile has a probability *J* of being recruited. During reproduction, individuals are assumed to contribute to the gamete pool in proportion to their size (the larger an individual, the larger its contribution to the gamete pool), and to self-fertilise at a fixed rate *α*.

**Figure 1:**
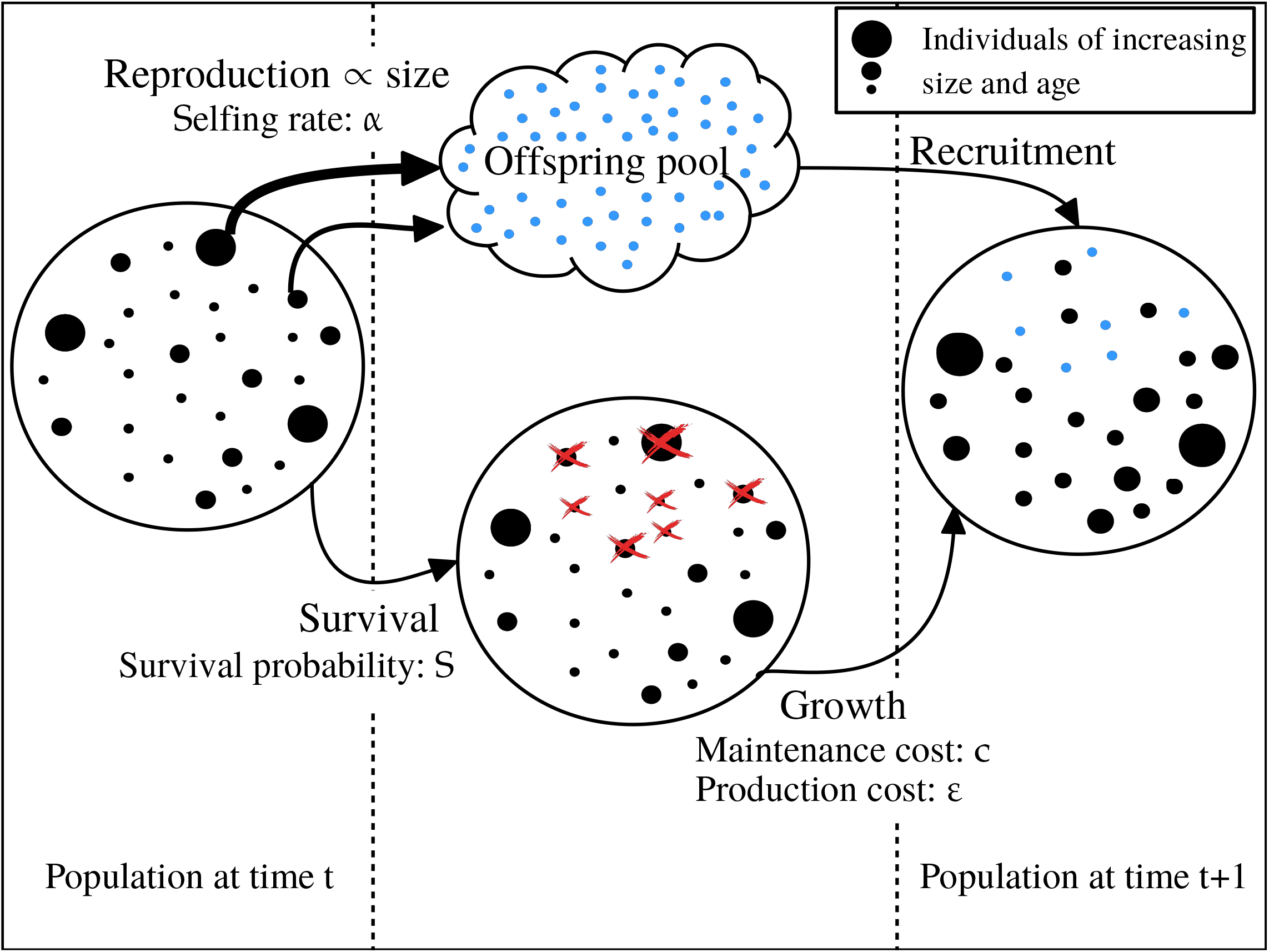
Demographic events over the course of one timestep. Deceased individuals are marked by a red cross, juveniles are depicted in light blue. Larger dots depict larger and older individuals.

### Growth model

We consider the growth model developed by West et al. (2001). The energy available for growth and maintenance at age *t*, *B_t_*, is assumed to scale as a ^3^/_4_-power law of body size as a result of allometry (Peters, 1983), so that

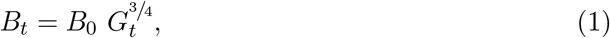

where *B*_0_ is the basal metabolic rate and *G_t_* is body size at age *t*. This energy can be subdivided into the energy required to maintain the existing body, controlled by a maintenance cost *c*, and the energy available to produce new body parts, controlled by a production cost *ε*, so that growth is fully described by the following differential equation

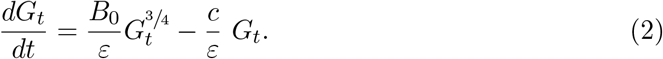

Under this model, individual size naturally saturates when the energy required to maintain the existing body equals the available energy (Fig. 2a-2b). Further details regarding the growth model are given in Appendix I.

**Figure 2:**
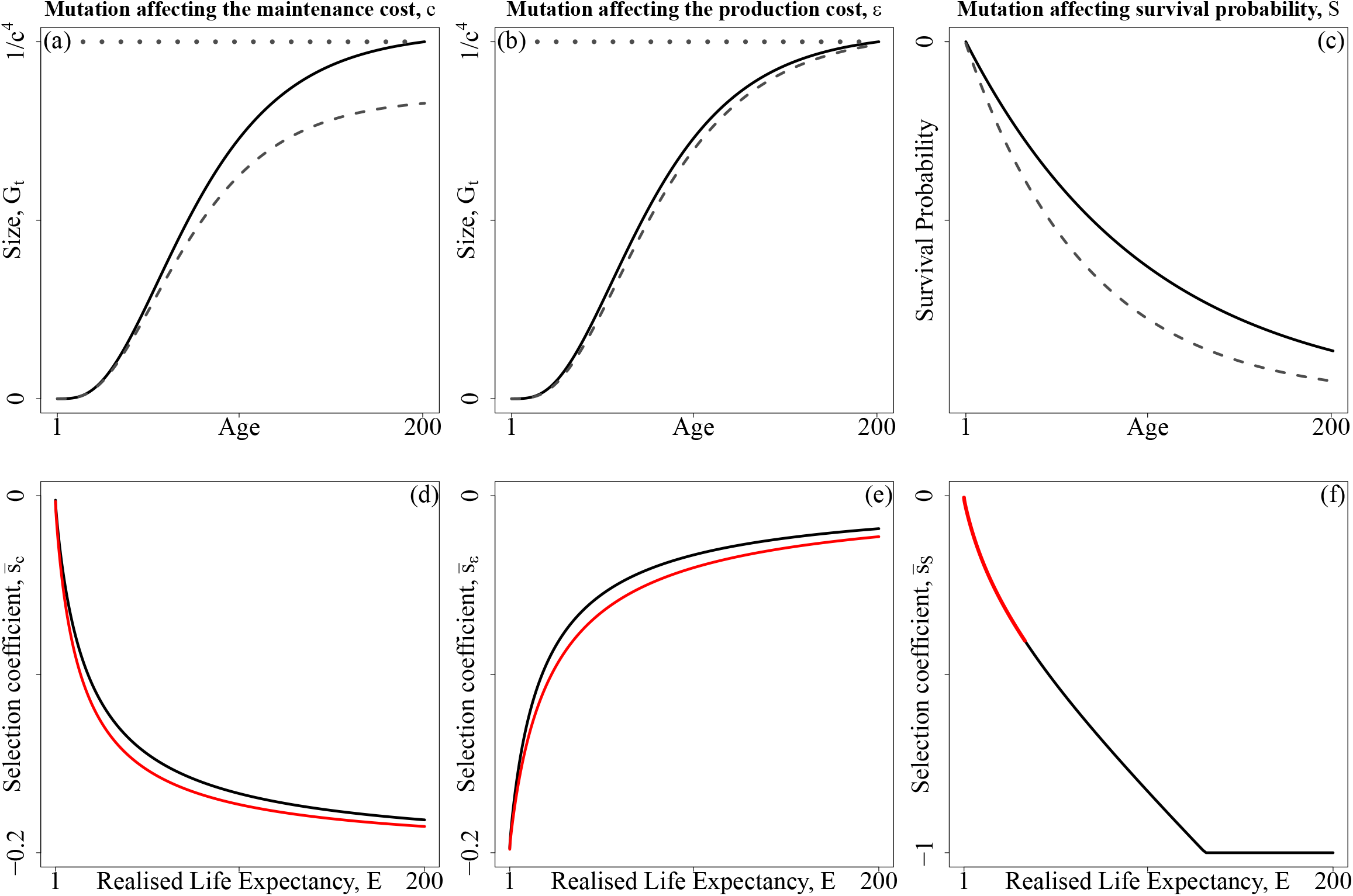
Phenotypic and leading order fitness effect of a mutation. Top row: phenotypic effects on growth or survival are presented as a function of age, the dashed line depicts the mutant phenotype while the solid line depicts the wild type (*c* = 0.001, *ε* = 0.01 and *s* = 0.05 for mutations affecting growth, and *S* = 0.99, *s* = 0.005 for mutations affecting survival). The dotted gray lines depict maximal size. Bottom row: Resulting effects of mutations on lifetime fitness as a function of life expectancy in a single locus case (black lines, *n* = 0), and 10 mutations segregate at other loci (red lines, *n* = 10). Note that the phenotypic effect of mutations (*s*) differs between mutations affecting growth and survival.

### Genetic assumptions

Mutations are assumed to occur at rate *U* (per haploid genome) at a large number of loci, which recombine at constant rate 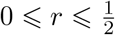. In three separate models, we consider mutations affecting three different traits, that is we consider mutations affecting one trait at a time (no pleiotropy). Mutations may affect growth by increasing either their bearer’s maintenance cost (*c*) or production cost (*ε*), or they may affect its survival. When mutations affect survival, they are assumed to decrease both their bearer’s probability of being recruited as a juvenile (*J*) and its adult survival probability (*S*). The effect of mutations is denoted *s*, with a dominance coefficient 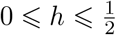. Loci affect traits multiplicatively, so that for any trait *z* (*z* ∈ {*c*, *ε*, *S*}), we have

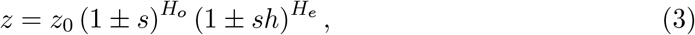

where *H_o_* (resp. *H_e_*) is the number of homozygous (resp. heterozygous) mutations born by the considered individual. The ± sign is used a general notation to indicate the fact that depending on the trait, deleterious mutations may increase it, as it is the case for the maintenance and the production cost; or decrease it, as it is the case for survival.

### Approximation of the expected number of mutations, inbreeding depression and mutation load

Our analytical model is built using the framework introduced by Barton and Turelli (1991) and generalized in Kirkpatrick et al. (2002). In this framework, the genetic dynamic of the population is described using allelic frequencies and genetic associations instead of genotypic frequencies. To do so, we define *X_i_* and 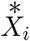 as the indicator variables associated with the paternally and maternally inherited allele at the *i^th^* locus, respectively. These variables are worth 1 if the deleterious allele is present at the considered position, and 0 otherwise. Thus, Equation (3), which depicts the phenotypic effect of mutations, can be rewritten in terms of these variables as

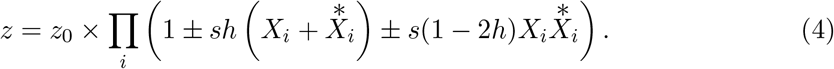

Indicator variables can in turn be used to define centered variables *ζ_i_* and 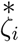, which are given by

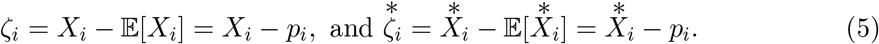

By definition, we have 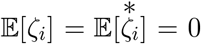, and expectations of products of these variables can be used to quantify how the population deviates from Hardy-Weinberg’s equilibrium, that is genetic associations in the sense of Barton and Turelli (1991), such as excesses in homozygotes, or linkage and identity disequilibria between loci for instance (see Appendix II.1 for details).

In order to obtain an analytical prediction of the average number of mutations per haploid genome at mutation-selection equilibrium, one needs to find the point at which the rate at which mutations enter the population through mutation during meiosis equals the rate at which mutations are removed from the population by selection. This can be done by following the change in allelic frequencies at selected loci between two timesteps, Δ*p_i_*, and looking for the point at which we have Δ*p_i_* = 0. We do so using two different approaches. In the first approach, we make the assumption that selective pressures acting on mutations at various life-stages can be summarised into a single lifetime fitness expression, so that the population can be studied as an adequately rescaled annual population (the Lifetime Fitness approach, LF - Appendix III.1 and Fig. S1). For each type of mutation, this approach allows us to gain insights into the way selection acts on mutations, by summarising it into a single lifetime selection coefficient, 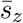, where *z* = *c*, *ε* or *S* depending on the considered trait (see Appendix III.1.1 for details). Besides, reasoning in terms of lifetime fitness is paramount to compute key quantities such as inbreeding depression or the mutation load. However, the LF approach fails to account for genetic associations correctly because it assumes that all selection occurs before reproduction at each timestep (see Fig. S1), so that it overlooks the fact that selection may occur both in parents and juveniles, that is before and after reproduction, respectively, which induces different effects of selection on genetic associations in these two subpopulations. Thus, in order to obtain approximations of the expected number of mutations per haploid genome accounting for genetic associations, we also use a second approach where we study each step of the life cycle successively (the Life Cycle approach, LC - Appendix III.2 and Fig. S2). We assume that the phenotypic effect of mutations is weak and that the number of segregating mutations is large, following the work of Roze (2015) that we adapted to the case of many mutations affecting a trait rather than fitness directly, in an age- and size-structured population.

### Simulations

In order to assess the validity of our analytical approximations, individual-centered simulations were run assuming diploid individuals depicted by two linear chromosomes of length *λ* (in cM), along which mutations occur stochastically during meiosis (Roze and Michod, 2010). The number of mutations occurring is sampled from a Poisson distribution with mean *U*, and their position on chromosomes are sampled from a uniform distribution. Recombination is modeled by exchanging segments between chromosomes. Similar to mutations, the number of crossing-overs is sampled from a Poisson distribution with mean *λ*, while their positions along chromosomes are randomly drawn in a uniform distribution. The population has a constant size *N*. At each timestep, an individual survives with probability *S*, which can depend on its genotype in the case of mutations affecting survival. If the individual does survive, it grows deterministically depending on its age and individual physiological growth costs (Fig. 2, Equation A5). If it does not, it is replaced by an offspring generated from the parental population, which includes the dead individual, and in which parents are chosen with a probability proportional to their size. The offspring is produced by self-fertilisation with probability *α*, and by random mating otherwise. We measure the average of the trait affected by mutations, the average number of mutations per haploid genome in the population, and inbreeding depression. Inbreeding depression is measured as the relative difference in lifetime reproductive success of selfed and outcrossed individuals. Individuals lifetime reproductive success is obtained by counting the number of times they are chosen as parents before they die. The mutation load in the sense of Crow (1958), that is the decrease in mean fitness of the population compared with a population with no mutations, is measured using the population average of the trait affected by mutations. This average is used to compute the expected mean lifetime fitness in the population (Equation A35). Then, this quantity is compared to the expected mean lifetime fitness when mutations are absent. All programs are available from GitHub (links are available from the journal office).

## 3 Results

In what follows, results will be presented as functions of the life expectancy of the population, *E*, that is the average number of mating events an individual is expected to live through before dying. Since survival is not assumed to vary with respect to age nor size, the probability of an average individual to survive up to exactly age *k* then die follows a geometric distribution of parameter 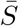, where 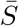 is the mean survival probability in the population, which is simply 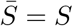 when mutations affect growth, and depends on mutations segregating in the population when said mutations affect survival. Thus, we have

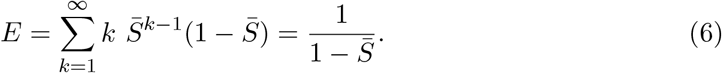

### 3.1 Mutation-selection equilibrium

To study the change in allelic frequencies at selected loci, one needs to quantify the intensity of selection acting on mutations. Generally speaking, the intensity of selection depends on how mutations modify the fitness of their bearer, and how the fitness of their bearer then compares with the fitnesses of all the individuals they are competing with, that is the distribution of fitness in the population. Hence, the intensity of selection faced by a mutation segregating at a given locus may depend on the presence of mutations at other loci, since said mutations may alter both the distribution of fitness in the population and how a mutation at the considered locus affects its bearer’s fitness. This second aspect is referred to as epistasis. There are numerous ways by which epistasis may be generated between loci (Phillips, 2008), which can roughly be split into two categories: genotype-to-phenotype epistasis on the one hand, and phenotype-to-fitness epistasis on the other hand. Genotype-to-phenotype epistasis refers to loci which influence each other’s expression in the determination of a phenotype, say genes involved in a common metabolic pathway, while phenotype-to-fitness epistasis refers to situations where loci, even if they influence phenotype independently, influence each other because the way the phenotype affects fitness is non-linear.

In our model, we assume that mutations affect a phenotype multiplicatively, so that genotype-to-phenotype epistasis does not occur. On the other hand, we do not impose any constraint on the shape of the fitness landscape. Instead, we let the genotype-to-phenotype-to-fitness map arise from biological assumptions. The resulting fitness landscape is non-linear, convex, and has singularities at the optima (Equation A35 in Appendix III.1, Fig. S3). Thus, phenotype-to-fitness epistatic interactions are generated, and have to be accounted for to obtain accurate mutation-selection balance approximations, so that quantifying the intensity of selection acting on a mutation at a given locus requires accounting for the average number of mutations it may be in epistatic interaction with. This can be done by following the change in the trait average as mutations segregate (Equation 7, see Appendices III.1.2 and III.2.2 for the derivation of this result under the Lifetime Fitness and the Life Cycle approach, respectively).

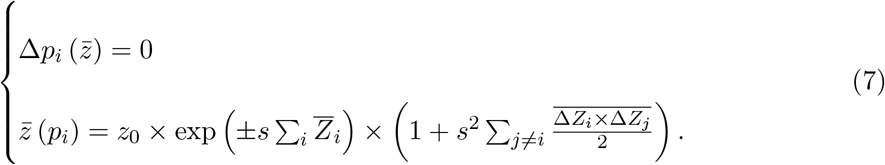

In the first line of Equation (7), Δ*p_i_* is the change in the frequency of the mutant allele at the *i^th^* locus, which depends on the trait average 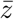. On the second line, 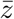 is the average of the trait in the population, which depends on the number of mutation per haploid genome ∑*_i_p_i_*. On this line, the ± sign indicates the fact that deleterious mutations may increase the average trait of the population, as it is the case for the maintenance and the production cost, or decrease it, as it is the case for survival. The term 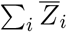 quantifies the effect on the trait of the ∑*_i_p_i_* mutations per haploid genome borne on average by individuals, neglecting the effects of genetic associations between loci, while the term 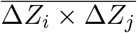 quantifies the effect of pairwise associations between loci on the trait average. Associations of higher order are neglected. Solving Equation (7) for *p_i_* and 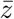 allows us to obtain predictions for the average of the trait and for the average number of mutations per haploid genome maintained at mutation-selection equilibrium, which we then use to compute the inbreeding depression and mutation load expected at equilibrium.

### 3.2 Fitness effect of mutations neglecting genetic associations

Under the Lifetime fitness approach, to leading order in *s* so that genetic associations between loci can be neglected, Equation (7) simplifies into

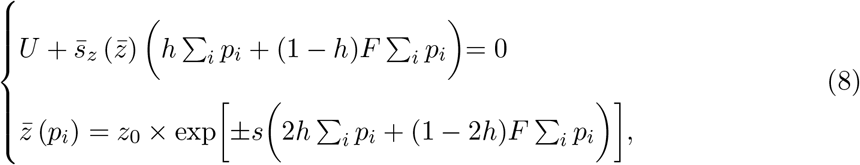

where 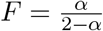 is the inbreeding coefficient. The derivation of this result is explained in Appendix III.1.2 (Equation A44). The first line of Equation (8) shows that when we neglect genetic associations between loci, the selective pressures acting on mutations are encapsulated in a single lifetime selection coefficient 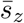, which is specific to each model and emerges from the application of the Taylor expansion methods presented in Appendix II.2.1 to lifetime fitness. Its expression for each type of mutation studied is given in Appendix III.1.1.

The phenotypic effect of each type of mutation (Fig. 2, top row), and the resulting leading order lifetime selection coefficients are presented in Fig. 2 (bottom row). These coefficients are all negative because we consider deleterious mutations, and depend on the population average of the trait, owing to the epistatic mechanisms described in the former section. Although it is not possible to fully disentangle the effects of epistasis from the effects of direct selection at the locus, one may gain insight into the sign of epistasis by studying the direction in which 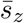 coefficients are changed when the number of segregating mutations increases in the population. The 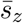 coefficients plotted in black were obtained assuming mutations are absent (*i.e*. 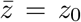), while those plotted in red were obtained assuming *n* = ∑*_i_p_i_* = 10 mutations were segregating at surrounding loci. In other words, the coefficients plotted in black in Fig. 2 (bottom row) represent the intensity of selection a mutation would face if a single locus was modeled, while the coefficients plotted in red depict the intensity of selection a mutation would face when in interaction with ten mutations at other loci.

Mutations affecting the maintenance cost *c* cause size differences to increase as individuals age (Fig. 2a). Hence, their fitness effect increases with life expectancy (Fig. 2d). Furthermore, epistatic interactions cause selection against mutations to increase that is, we observe negative epistasis. On the contrary, mutations affecting the production cost *ε* do not affect individuals’ maximal size, as they asymptotically tend to the same size, but rather the speed at which they reach it (Fig. 2b). Therefore, the growth delay mutated individuals accumulate in early years fades away in older individuals. This causes selection against mutations affecting *ε* to decrease with respect to life expectancy, because they become gradually neutral in older age-classes. However, epistasis also causes selection against mutations to increase, so that it is negative in this case as well.

Mutations affecting survival cause individuals to perform less mating events in a life-time. They also cause mutated individuals to perform less well during mating events, because they tend to be younger than unmutated individuals (Fig. 2c). Thus, age-structure increases selection against mutations affecting survival. Furthermore, selection against mutations affecting survival strongly increases as life expectancy increases (Fig. 2f), contrary to mutations affecting growth costs, whose fitness effects remain moderate (Fig. 2d-e). Furthermore, strong positive epistasis is observed in the case of mutations affecting survival, as said mutations decrease the life expectancy of the population, so that the intensity of selection against mutations sharply decreases when the number of segregating mutations increases.

Overall, the results described in Fig. 2 show that mutations affecting different traits on the same genotype-to-phenotype-to-fitness map face very different selective pressures, both in magnitude and in the way they vary with life expectancy.

### 3.3 Average number of mutations, inbreeding depression and mutation load

The intensity of selection acting on mutations does not depend on the values of *c* and *ε*, but on the ratio 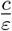 (Appendix III.1.1). Thus, in what follows, results are described using this ratio. Figure 3 presents results for the average number of mutations per haploid genome maintained at equilibrium, and the resulting inbreeding depression and mutation load for *h* = 0.25 and 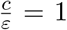 (other parameter sets are shown in appendix as results are qualitatively similar, Fig. S5 to S12). Analytical predictions, which are depicted by solid lines, are obtained by solving Equation (7) numerically using the LC approach (Appendix III.2) and accounting for pairwise genetic associations, and dots depict simulations results.

**Figure 3:**
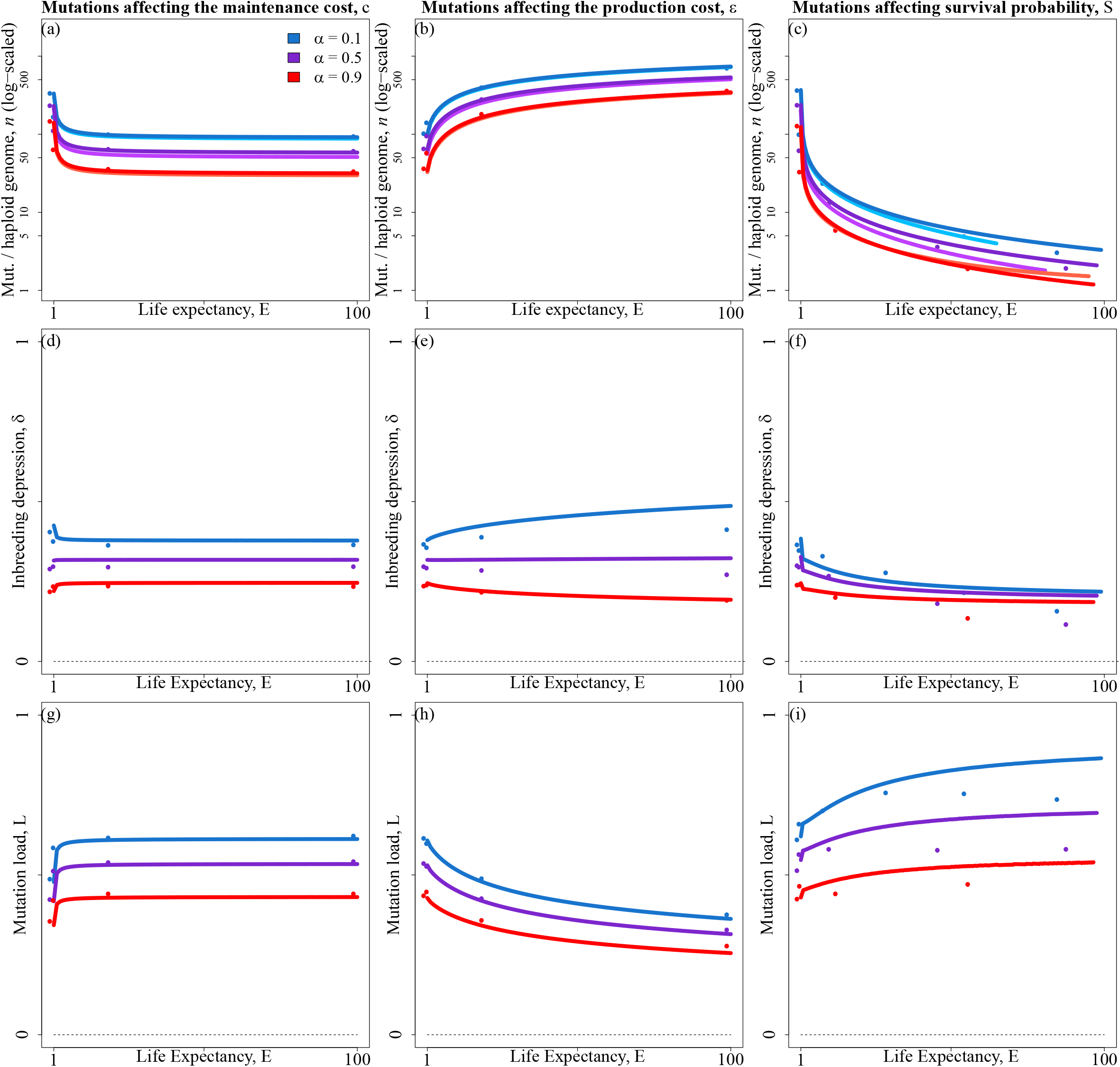
Average number of mutations per haploid genome (*n*, top row), inbreeding depression (*δ*, middle row), and mutation load (*L*, bottom row) as a function of life expectancy (*E*), for three selfing rates : *α* = 0.1 (blue), *α* = 0.5 (purple) and *α* = 0.9 (red). Each column corresponds to one type of mutation. Dots: simulation results. Lines: analytical predictions accounting for genetic associations between loci using the LC approach. Parameters shown here are 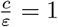, *U* = 0.5, *s* = 0.005, *h* = 0.25.

**Figure 4:**
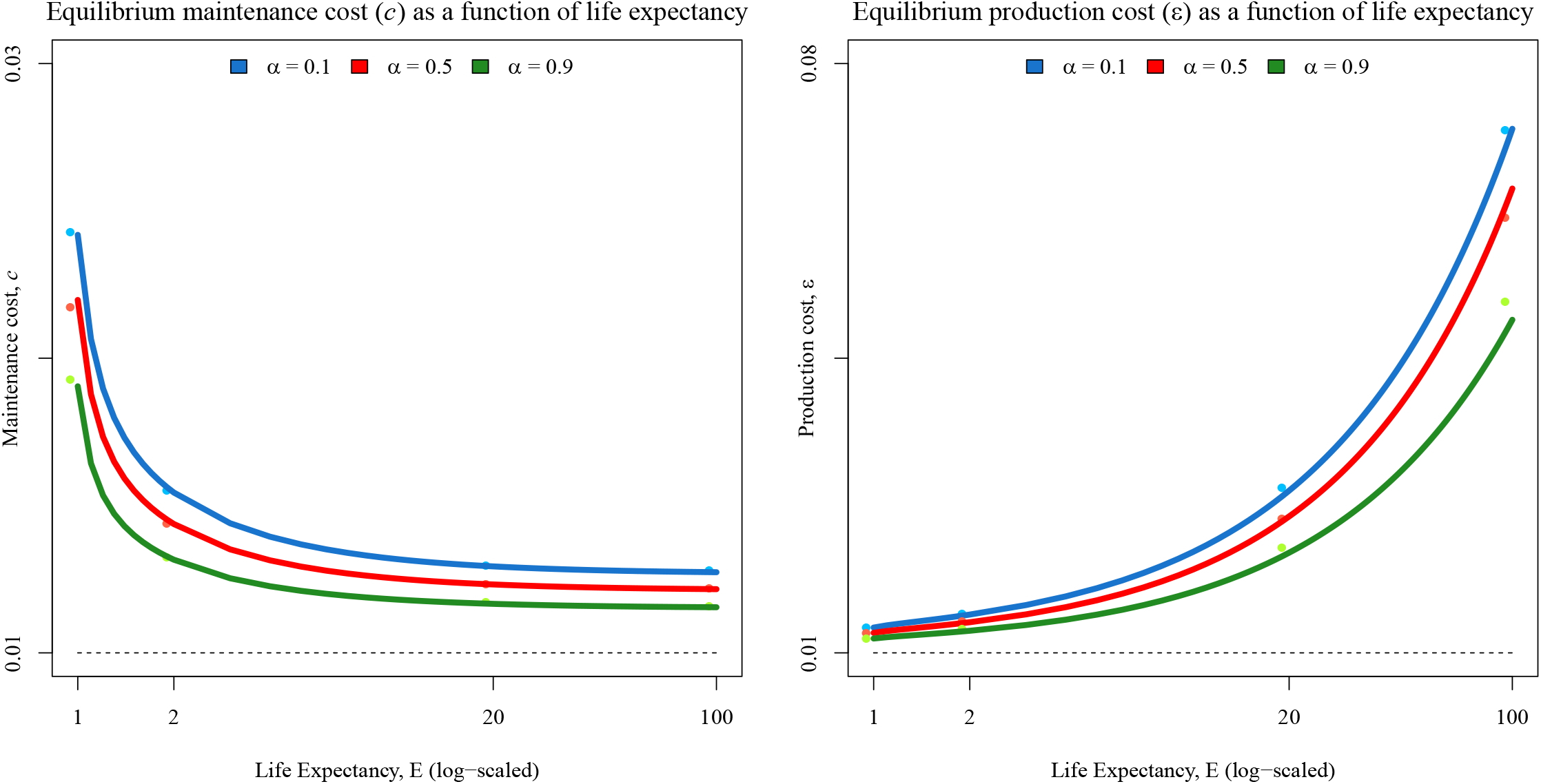
Equilibrium maintenance (left) and production (right) costs as function of life expectancy (log-scaled), for various selfing rates. Dots depict simulation results while lines depict analytical predictions. Parameters shown here are *c* = *ε* = 0.01, *h* = 0.25, *s* = 0.005.

The first row in Fig. 3 presents the log-scaled average number of mutations per haploid genome maintained at mutation-selection balance. For mutations affecting the maintenance cost and survival, the average number of mutations per haploid genome maintained at equilibrium (*n* = ∑*_i_p_i_*) decreases as life expectancy increases (Fig. 3a and 3c), because selection against mutations increases. This effect is more marked for mutations affecting survival because selection is stronger in this case (Fig. 2d and 2f). Conversely, *n* increases as life expectancy increases for mutations affecting the production cost, because selection against these mutations weakens as life expectancy increases (Fig. 2e). In every case, *n* decreases as the selfing rate increases due to the purging effect of self-fertilisation.

The middle row in Fig. 3 shows the levels of inbreeding depression generated by mutations maintained at mutation-selection balance. Large differences in the number of deleterious mutations maintained per genome do not translate into strong variations in inbreeding depression with respect to life expectancy. However, variations are still observed and differ between mutation types. Indeed, inbreeding depression always decreases with life expectancy when mutations affect survival, irrespective of the selfing rate (Fig. 3f), contrary to mutations affecting growth. Mutations affecting *c* generate higher inbreeding depression in more short-lived species at low selfing rate, but this pattern is reversed at higher selfing rates (Fig. 3d). Conversely, mutations affecting *ε* generate higher inbreeding depression in more long-lived species for low *α* and this pattern is reversed for high *α* (Fig. 3e). This is the result of the interaction between life expectancy, the shape of the fitness landscape and selfing, which is difficult to disentangle as these elements interact in non-trivial ways. Indeed, the shape of the fitness landscape changes with respect to life expectancy, which causes variations in the intensity of selection and epistatic interactions, and selfing changes the distribution of fitness by altering homozygosity and by promoting the purging deleterious mutations in doing so, so that it too alters epistatic interactions.

The bottom row in Fig. 3 presents the mutation load measured in the sense of Crow (1958) which results from the mutations segregating at mutation-selection balance. Intuitively, one would assume that having more deleterious mutations segregating in the population should lead to a higher mutation load. Yet, the mutation load is lower when mutations are more numerous for all three mutation types in our model. Importantly, this result is not a mere reflection of differences in the absolute strength of selection acting on mutations in populations with different life expectancies. Otherwise, changing the phenotypic effect of mutations in a given model and for a given parameters set would also change the mean phenotypic deviation from the optimum, and therefore the mutation load. This is not what we observe. Indeed, Figure S4 in Appendix V shows that when the effect of mutations is made ten times larger, this does not cause differences in the equilibrium phenotypic deviation, although the number of segregating mutations at equilibrium was considerably lower (because selection was stronger). This observation is consistent with results obtained by previous authors (Bataillon and Kirkpatrick, 2000), who showed that when the population exceeds a particular size, the mutation load becomes independent from the strength of selection acting on mutations. We argue that the differences we observe in terms of mutation load are imputable to differences in the shape of the fitness landscapes (Fig. S3), which are not fully captured by differences in the efficacy of selection.

### 3.4 Consequences for life-history evolution

The mutations segregating in the population cause the mean phenotype to deviate from its initial value. As the intensity of selection acting on mutations depends on this mean phenotype, this leads to a joint equilibrium for both the mean phenotype and the number of mutations segregating in the population (Appendices II and III). This equilibrium varies with respect to life expectancy and mating system. Increasing the selfing rate slightly decreases the deviation of the mean phenotype from its initial value, because selfing induces a better purging of mutations. More importantly, the equilibrium phenotypes vary significantly between populations with different life expectancies. Indeed, as species become more long-lived, the equilibrium maintenance cost decreases, so that maximal size increases, and the equilibrium production cost increases, so that growth is slowed down. This means that if, for any reason, life expectancy changes in a species, the growth strategy should also be changed as a consequence of the selective pressures acting on deleterious mutations maintained at mutation-selection balance being altered.

## 4 Discussion

### Inbreeding depression

In this paper, we set off to study the magnitude and variation of inbreeding depression generated by mutations affecting survival or growth in relation to life expectancy. We showed that inbreeding depression at mutation-selection balance varies weakly with respect to life expectancy when mutations affect growth, while inbreeding depression decreases more sharply as life expectancy increases in the case of mutations affecting survival. We showed that these differences between mutation types can be attributed to differences in the intensity of selection and how it varies with respect to life expectancy. In any case, our results only agree with empirical data in a limited region of the parameters space, that is when selfing is rare and selective pressures acting on mutations decrease as life expectancy increases, as it is the case for mutations affecting the production cost most noticeably. Although these results may at first glance seem to point to a more preponderant role for somatic mutations accumulation in generating the sharp increase in inbreeding depression observed in more long-lived species, we argue in what follows that they may rather point us towards the type of traits mutations should affect in order to produce this empirically observed pattern, irrespective of the way mutations are produced.

As stated above, differences in the magnitude of inbreeding depression with respect to life expectancy were quantitatively small in every investigated case when mutations affected growth, that is either the maintenance or the production cost, even for low dominance coefficients and high mutation rates. This result can be explained by the moderate variations in the intensity of selection on growth-related traits with respect to life expectancy, and by the fact that we only considered mutations with weak phenotypic effects, which in this case translated into weak fitness effects. Therefore, this result is consistent with previous population genetics work that showed that inbreeding depression becomes independent of the intensity of selection when selection is weak and the population is large (e.g. Charlesworth et al., 1990; Bataillon and Kirkpatrick, 2000; Roze, 2015). As for mutations affecting survival, their dynamics were studied by Morgan (2001) prior to this study in a perennial, but not age-structured population. They concluded that inbreeding depression should quickly decrease as life expectancy increases, and that significant outbreeding depression should be observed in long-lived species. They argued this result could be attributed to the greater variance in fitness observed among the offspring produced by self-fertilisation, which led to a higher mean lifetime fitness among them. Although they did not state it explicitly, this result stems from the fact that a very small number of mutations were maintained at mutation-selection balance in long-lived species under the parameter sets they investigated. Indeed, Morgan (2001) considered large phenotypic effects of mutations (*s* = 0.1 and *s* = 1.0), which resulted in tremendous lifetime fitness effects and therefore in the maintenance of almost no mutation at equilibrium. This led variance in fitness to dominate over other aspects. Here, we investigated mutations with much lower phenotypic effects (*s* = 0.005), and did not observe the same patterns. In fact, we show that although inbreeding depression still decreases with life expectancy, differences are considerably reduced, and outbreeding depression is no longer observed. Thus, if substantial inbreeding depression is observed on traits related to survival in long-lived species, we conclude that this should be caused by mutations with very weak phenotypic effects, or whose effects are limited to early stages of life, as mutations would otherwise not be maintained in the population.

Taken together, our results and those of Morgan (2001) show that variations in the fitness effect of mutations with respect to life expectancy can generate differences in inbreeding depression, provided that these variations are strong enough. Furthermore, we showed that inbreeding depression maintained at mutation-selection balance decreases when the fitness effect of mutations increases, as said mutations are more efficiently purged from the population. This implies that mutations facing sufficiently strong selective pressures, which decrease sharply as life expectancy increases, could in principle generate the increase in inbreeding depression observed in more long-lived species. In the light of this result, we propose that asking on what traits selection is expected to be strong and to decrease significantly as life expectancy increases in plants is a track worth following in order to further our understanding of the mechanisms underlying inbreeding depression. Importantly, this result is not in contradiction with the somatic mutations accumulation hypothesis formulated by Scofield and Schultz (2006). Indeed, the intensity of selection acting on mutations matters for their persistence in the population and for the magnitude of inbreeding depression generated by them, regardless of the way mutations are produced.

### Mutation load

Contrary to what one might intuitively expect, we showed that more mutations maintained at mutation-selection balance led to a lower mutation load in all three of our models, and that this result is not a straigthforward consequence of differences in the intensity of selection acting on mutations, in agreement with results obtained by previous authors (Bataillon and Kirkpatrick, 2000). Instead, it is a consequence of differences in the shape of fitness landscapes between species with different life expectancies on the one hand, and of the interaction between the mean phenotype and the intensity of selection on the other hand. This result highlights the fact that incorporating a phenotypic dimension to population genetics studies may lead to counter-intuitive results. In particular, when fitness landscapes are obtained as the result of biological assumptions, and not arbitrarily assumed to be of a particular shape, unusual interactions with various aspects of species life-histories may result in novel predictions. For instance, in the present study, we showed that the mutation load may behave very differently depending on the trait affected by mutations and life expectancy. Therefore, we conclude that comparing mutation loads between species with contrasting life-histories may sometimes be misleading, and would likely require trait-specific approaches.

### Life-history evolution

Plants vary widely in life expectancy and stature, with life expectancies ranging from a few weeks to hundreds, possibly thousands of years, and stature spanning several orders of magnitude across Tracheophytes (Ehrlén and Lehtilä, 2002). These variations are correlated. Indeed, long-lived species tend to grow slower than short-lived species (Salguero-Gómez et al., 2016). In life-history traits evolution theory, this type of correlation is usually interpreted in terms of trade-offs, with populations evolving towards the evolutionarily stable allocation of resources between growth, survival and reproduction, given a number of constraints (Stearns, 1992). In this paper, we have shown that the equilibrium maintenance and production costs differ between life expectancies, causing more long-lived species to grow slower but ultimately larger than more short-lived species, as commonly expected. However, the mechanism underlying this result is completely different. Indeed, in our model, life-history traits do not coevolve in response to trade-offs. Instead, the equilibrium growth costs are modified when life expectancy varies because the selective pressures acting on the many mutations affecting these traits are altered, leading to a more or less efficient purging of said mutations and thereby phenotypic differences at mutation-selection equilibrium. Thus, our results suggest that life-history traits may sometimes coevolve regardless of trade-offs, because a change in a given trait may alter the efficiency of purging of deleterious mutations affecting other traits.

### Epistasis emerging from the fitness landscape

The phenotypic dimension of the genotype-to-phenotype-to-fitness map is usually overlooked in mutation load dynamics studies. Indeed, most theoretical investigations interested in such matters consider mutations affecting fitness multiplicatively (e.g. Kondrashov, 1985; Charlesworth et al., 1990; Roze, 2015), so that fitness is the phenotype affected by mutations and the phenotype-to-fitness map (the fitness landscape) is strictly linear. Hence, neither phenotype-to-fitness epistasis nor genotype-to-phenotype epistasis occur in these models. Some authors however considered the effects of epistasis on mutation load dynamics.

Charlesworth et al. (1991) studied uniformly deleterious mutations under synergistic selection. Because they assumed mutations to affect fitness directly too, no phenotype-to-fitness epistasis occured in their model. On the other hand, genotype-to-phenotype epistasis occured because they assumed mutations to affect fitness synergistically. Abu Awad and Roze (2019) generalised Charlesworth et al. (1991)’s results, showing that their model is equivalent to assuming fixed negative pairwise epistasis between loci. Furthermore, they showed that different forms of pairwise epistasis have different effects: while additive-by-additive epistasis lowers the frequency of mutations by increasing selection, additive-by-dominance and dominance-by-dominance epistasis tend to increase inbreeding depression by making homozygotes less fit.

Phenotype-to-fitness epistasis was also considered in the case of mutations affecting an abstract trait (or set of traits) under stabilising selection (e.g. Abu Awad and Roze, 2018, 2019). These studies assumed mutations to affect the trait(s) additively, so that no genotype-to-phenotype epistasis occured, but assumed the fitness landscape to be gaussian (or quasi-gaussian), so that phenotype-to-fitness epistasis occured as the landscape was non-linear. However, because the fitness landscape was symmetrical with respect to the optimal phenotype and mutations occured in both directions at the same rate, epistasis was null on average (Gros et al., 2009). Yet, it influenced mutation load dynamics through variance effects, by generating associations between alleles with compensatory effects, and decreasing the efficiency of selection on deleterious alleles when these associations stay moderate.

In this paper, we studied the dynamics of mutations affecting growth and survival using a physiological growth model (West et al., 2001). Hence, the fitness effect of mutations emerged from biological assumptions instead of being assumed a priori. The resulting fitness landscape had singularities at the optimal phenotypes, as in the absence of trade-offs, immortality (*S* = 1) is the optimal survival strategy and leads to an infinite life expectancy, and null growth costs (*c* = *ε* = 0) are the optimal growth strategy, which leads to infinitely large individuals. Furthermore, these optimal phenotypes were situated on the boundaries of the landscape for all three traits, as negative growth costs or survival probabilities greater than one are meaningless. As a consquence, mutations were always under directional selection in our model. We hence focused on the case of mutations increasing growth costs or decreasing survival, that is deleterious mutations, as opposite mutations would always be favoured and eventually reach fixation. Besides, the fitness landscape was non-linear (Fig. S3). Hence, although we assumed mutations to affect phenotypes multiplicatively, so that no genotype-to-phenotype epistasis occured, phenotype-to-fitness epistasis was generated. Contrary to models assuming stabilising selection (e.g. Abu Awad and Roze, 2018, 2019), epistasis was non-zero on average in our model, and its direction (positive or negative) differed depending on the phenotype affected by mutations. It was however difficult to isolate the effects of epistasis in our model because pairwise epistatic interactions are not sufficient to capture these effects, as considering only pairwise interactions is equivalent to linearising the effect of mutations on the trait average, which only works when very few mutations segregate in the population.

## APPENDIX

### I Growth model

At any age *t*, we assume that an individual is composed of *G_t_* identical units (say, branches or buds), and has a resting metabolic rate *B_t_*. This metabolic rate can be subdivided into energies spent on maintenance and growth as follows (West et al., 2001),

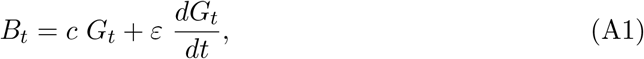

where *c* is the energy required to maintain a single unit, and *ε* is the energy required to create a new unit. Moreover, the resting metabolic rate can be expressed as

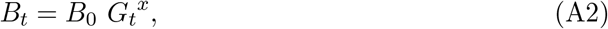

where *B*_0_ is the species’ basal metabolic rate and *x* is the allometric coefficient (Peters, 1983). In vascular plants and animals, the value *x* = ^3^/_4_ is generally used in growth models on the basis of both theoretical (e.g. West et al., 1997, 1999), and empirical arguments (e.g. Peters, 1983; Enquist et al., 1998; West et al., 2001), and will thus be used in what follows. Different exponent values would however likely yield qualitatively similar results, provided that they remain positive and smaller than one, because the rate at which the metabolic rate *B*_0_ *G_t_^x^* increases with respect to size would still slow down as size increases so that the energy available to the individual would at some point equal the energy required for its maintenance *c G_t_*, thereby reaching a plateau which is given by

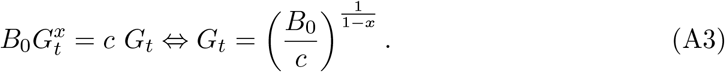

Using Equation (A2), Equation (A1) can be rearranged into the following differential equation

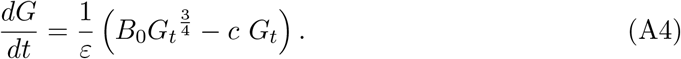

Solving Equation (A4) for *G_t_*, and setting *B*_0_ = 1 for convenience, we obtain

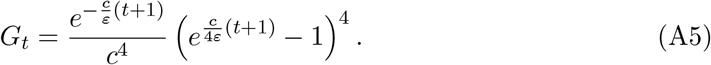

This growth function naturally saturates, when the energy required to maintain existing units becomes too large for new units to be produced. The size at which it saturates is given by

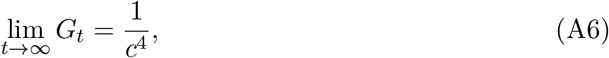

which is equal to the plateau given in Equation (A3), with *B*_0_ = 1 and 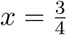.

### II General recursions for the effects of selection and reproduction under partial selfing

In this paper, we use two different approaches to study our model. The first approach makes the assumption that an age-structured population can be viewed as a rescaled annual population (the Lifetime Fitness approach, LF), so that individuals first undergo lifetime selection, then reproduce. The second method proceeds in a more detailed manner, by studying each step of the life cycle successively (the Life Cycle approach, LC). In both approaches, the objective is to derive an approximation for the change in frequency of deleterious alleles at the selected loci, Δ*p_i_*, in order to find the expected number of mutations maintained at mutation-selection balance in the population by solving Δ*p_i_* = 0.

Irrespective of the method, one may derive a general approximation of the relative fitness for an individual as a function of its genotype during any given selection stage. In this section, after introducing our theoretical framework (Barton and Turelli, 1991; Kirkpatrick et al., 2002), we derive such an approximation assuming fitness is a function of a trait *z*, and compute general recursions for the effects of selection on allelic frequencies and genetic associations. Then, we compute general recursions for the effects of reproduction.

#### II.1 Defining variables

Let *X_i_* and 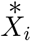 be the indicator variables associated with the paternally and maternally inherited alleles at the *i^th^* locus. These variables are worth 1 when allele *a* is present at the position they are associated with, and 0 otherwise. Let us also define the centered variables associated with the paternally and maternally inherited alleles at the *i^th^* locus, *ζ_i_* and 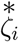, as

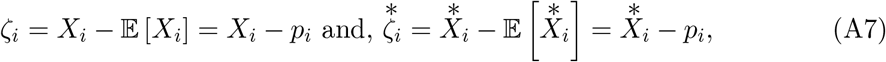

where 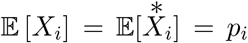 is allele *a*’s frequency at the *i^th^* locus. The expectation of products of these centered variables allows one to quantify genetic associations between positions in the genome, that is, these variables allow one to quantify to what extent the population deviates from Hardy-Weinberg’s equilibrium. For example, the expectation 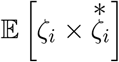 measures to what extent the homozygosity deviates from the panmictic expectation at locus *i*.

##### Condensed notations and simplification rule

To make recursions clearer, let us introduce the notation

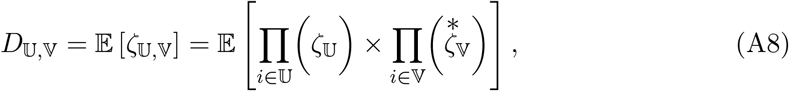

where 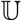 and 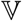 are sets of positions paternally and maternally inherited, respectively (in the previous example, 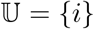 and 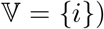, so that the excess in homozygotes at locus *i* is denoted *D_i,i_*. Sets 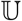 and 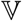 can contain positions situated at the same locus or at different loci, and can be empty. For instance, Association *D*_*ij*,∅_ which in terms of sets corresponds to 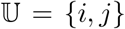, and 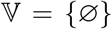, quantifies the linkage disequilibrium between alleles at loci *i* and *j* on the paternally inherited chromosome.

Besides, individuals are hermaphroditic and no sex-specific effect is assumed in our model, so that we always have

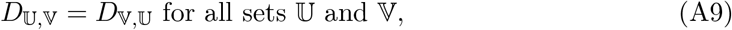

hence in our former example, we have *D*_*ij*,∅_ = *D*_∅,*ij*_. Again, to make recursions shorter, we will use the notation

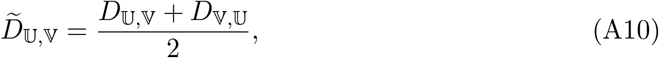

and when possible, we will omit empty sets. For example, we have

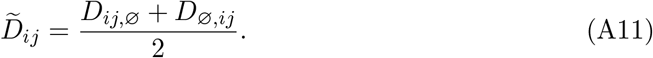

Finally, as shown in Kirkpatrick et al. (2002), repeated indexes sometimes appear in recursions, and can be dealt with using the relationship

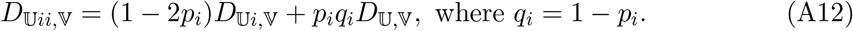

##### Genetic associations relevant to our model

When mating is not random, as it is the case in our model because of self-fertilisation, non-zero genetic associations are maintained between and within loci at equilibrium. Since we model an infinite number of loci, accounting for all associations arising between them is not possible as infinitely many combinations of genetic positions would have to be considered. Luckily, most genetic associations remain negligible at equilibrium when selection is weak, so that we may confine ourselves to taking into account within-locus and pairwise between-loci associations.

Furthermore, different kinds of genetic associations are affected by selection differently, so that further simplifications can be applied (Roze, 2015). Indeed, association 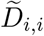, which depicts the excess in homozygotes at locus *i*, is generated by inbreeding even in the absence of selection, and direct selection acting on this association can be neglected when mutations segregating at other loci are numerous, because indirect selection becomes much stronger. Besides, 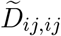, which reflects correlations in homozygosity between loci *i* and *j*, is also generated by inbreeding even under neutrality, and is little affected by selection, so that we can neglect the effects of selection on this association.

Other between-loci associations are generated both by inbreeding and selection. Indeed, 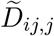 is generated by selection at locus *i* and is of order *s*, while 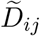 and 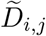 result from selection affecting both loci, and are of order *s*^2^. Following Roze (2015), we obtain leading order approximations in *s* for these associations, so that 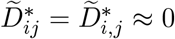 at quasilinkage equilibrium. Thus, in what follows, we will only be concerned with associations 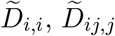 and 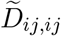, and neglect the rest of them.

#### II.2 Selection

##### II.2.1 Deriving relative fitness

For any trait *z*, assuming mutations affect this trait multiplicatively, an individual’s trait value can be written in terms of indicator variables as

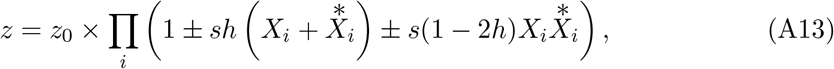

where ± is replaced with + or − depending on whether mutations are assumed to increase (e.g. physiological growth costs) or decrease (e.g. survival probability) the trait, *s* is the phenotypic effect of the mutant allele, and *h* is its dominance coefficient. The log-value of the trait is then

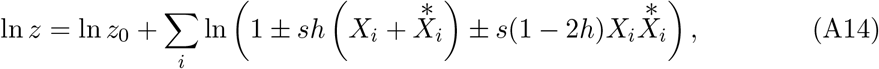

which, under the assumption that selection is weak, *i.e*. that *s* is small, can be approximated by

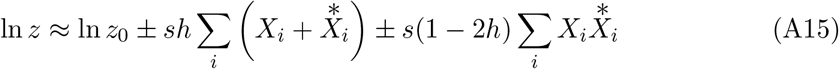

to leading order in *s*. Injecting *ζ*-variables into Equation (A15), it can then be rearranged into

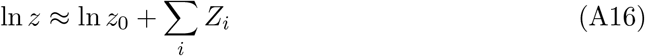

with 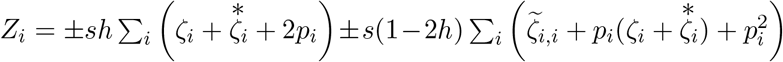. Hence, the mean log-value is given by

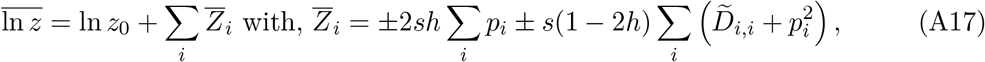

and one may express the trait value *z* as

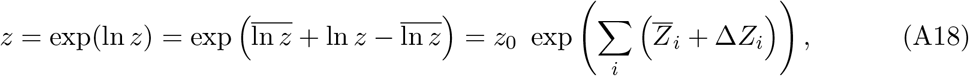

with 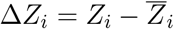. Therefore, the trait average, 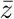, is simply

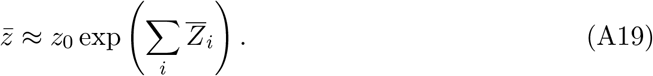

All else being fixed, an individual’s fitness during a given selection stage, *w*(*z*), is a function of its trait value *z*. The mean fitness is a constant, given by 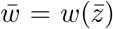, and the relative fitness 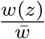, to second order in *s*, is given by

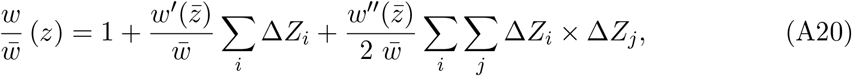

where *w*′ and *w*″ are the first and second derivatives of *w* with respect to Δ*Z_i_*. Neglecting terms in *i* = *j*, and denoting 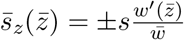 and 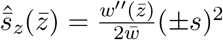, this is

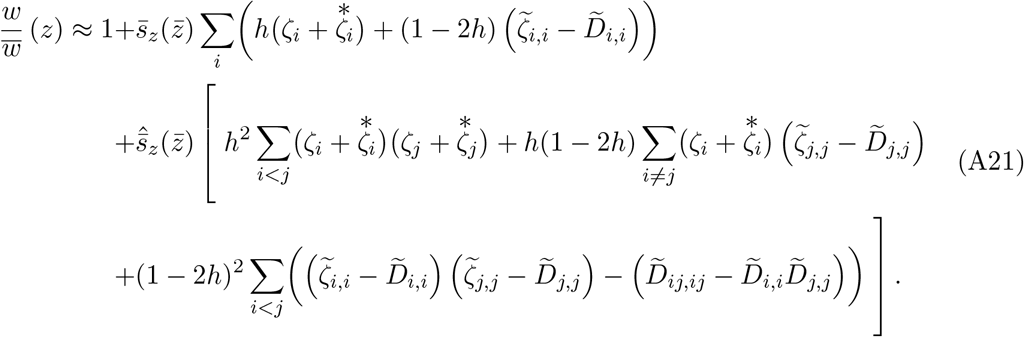

The leading order selection coefficient 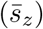 encapsulates the effects of selection acting directly on loci, while the second order selection coefficient 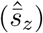 quantifies the effects of indirect selection between pairs of loci. We neglect higher order selection coefficients. Note that 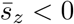 in all our models since we consider deleterious mutations.

##### II.2.2 Effects of selection of allelic frequencies and genetic associations

###### Allelic frequencies change due to selection

The change in allelic frequencies at the *i^th^* selected locus owing to selection at a given stage is

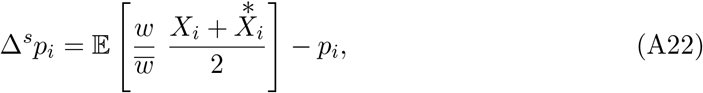

which, using Equation (A21), to second order in *s* and neglecting terms in 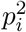, that is assuming mutations are rare, yields

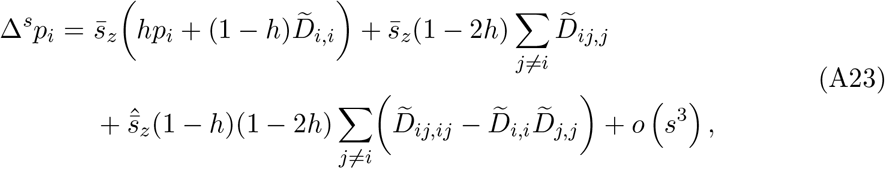

###### Effect of selection on associations between selected loci

We compute the effect of selection on genetic associations to leading order in *s* (first line in Equation A21). As shown by Roze (2015), the effects of selection on 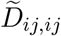 can be neglected and 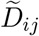 and 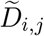 are of order *s*^2^. Hence, we only need to consider the effects of selection on 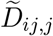 and 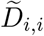, which can be computed using Equation (A24).

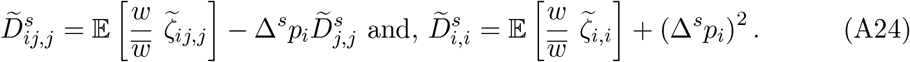

To leading order, Δ*^s^p_i_* simplifies to

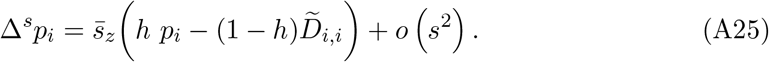

Hence, 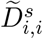 and 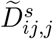 are given by

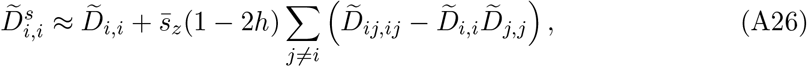

and,

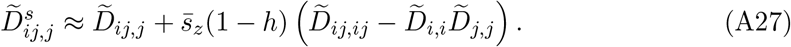

#### II.3 Reproduction and mutation under partial selfing

As we model infinitely many loci, we may assume that mutations never occur two times at the same locus. Since they are not affected by recombination and self-fertilisation, allelic frequencies following reproduction are simply given by

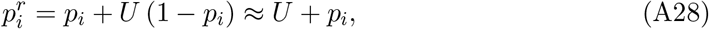

where *p_i_* here depicts the frequency of the deleterious allele at locus *i* just before reproduction. On the other hand, while we neglect the effects of mutation on genetic associations, they are affected by recombination and selfing. Following reproduction, denoting 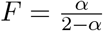, we have

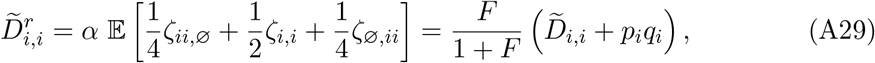

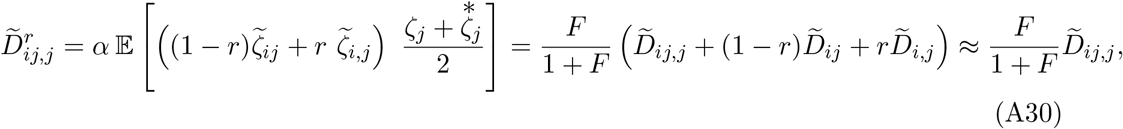

and,

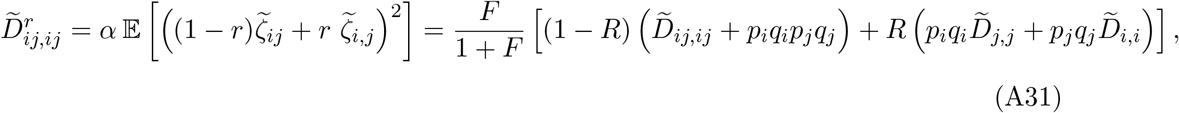

with *R* = 2*r*(1 – *r*) and where 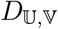 associations and allelic frequencies *p_i_* are measured just before reproduction.

### III Analytical results

In Appendix II, we derived general recursions for the effects of reproduction (involving mutation, non-random mating and recombination) and selection at any given selection stage. In this section, our aim is to apply these general recursions to our different models and analytical approaches. For each analytical approach, we begin with deriving the leading and second order selection coefficients arising from the models, then present analytical results at quasi-linkage equilibrium and explain how they are used to obtain an approximation for the average number of mutations per haploid genome.

#### III.1 Lifetime fitness approach

In the LF approach, we make the assumption that all the selective pressures acting on mutations can be summarised into a single lifetime fitness expression, so that the population can be studied as a rescaled annual population. Thus, in this analytical model, timesteps begin with lifetime selection, and are followed by reproduction (Fig. S1).

**Figure S1:**
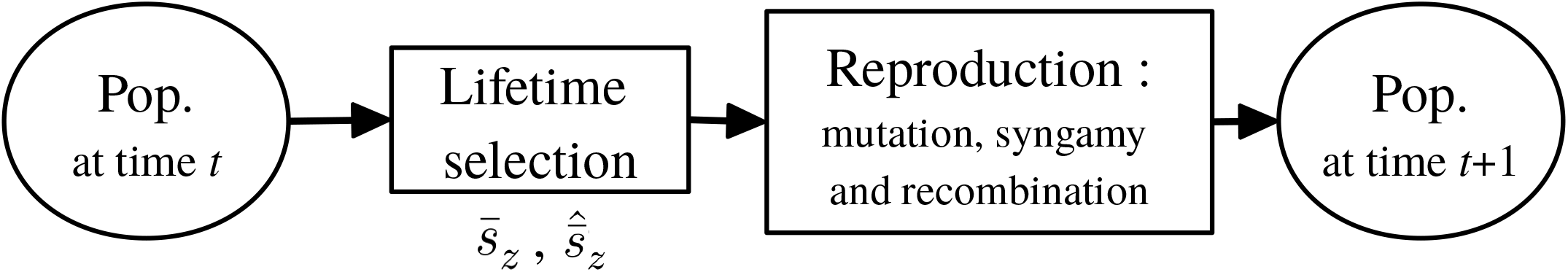
Steps followed over the course of one timestep with the LF approach

##### III.1.1 Lifetime fitness and lifetime selection coefficients

In this section, we compute the lifetime reproductive output of an individual given its phenotype, which is used as a measure of lifetime fitness, and derive the leading and second order lifetime selection coefficients resulting from this expression.

###### Lifetime fitness

The size of an individual at age *t* (*Gt*) is the solution of Equation (2). Given the individual’s maintenance and production costs, *c* and *ε*, it is

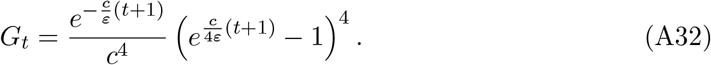

Hence, the contribution to reproduction of an individual up to age *τ*, 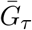, is given by

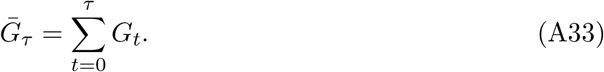

Furthermore, the probability that, given its survival probability *S*, an individual lives up to exactly age *τ* then dies 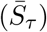 is

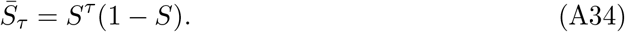

Hence, the lifetime fitness of an individual (*W*) is given by

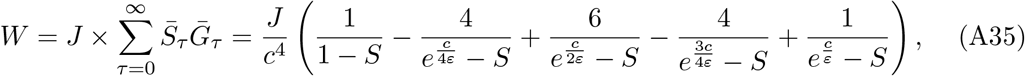

where *J* is the recruitment probability of the individual as a juvenile.

###### Selection coefficients

Let us now write down the lifetime selection coefficients obtained with the method described in Appendix II.2. For any trait *z*, these coefficients are obtained by applying Equation (A20) to Equation (A35). We show that they depend on the ratio 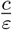, rather than *c* and *ε* absolute values. In what follows, we set *J* = 1 for convenience.

###### Mutations altering *c*

The leading order lifetime selection coefficient for mutations altering *c* is given by

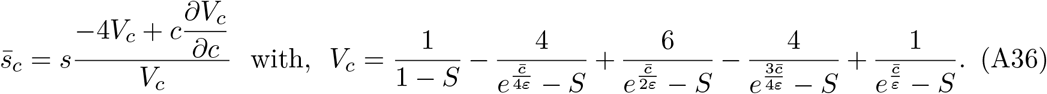

As for the second order lifetime selection coefficient, 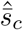, it is given by

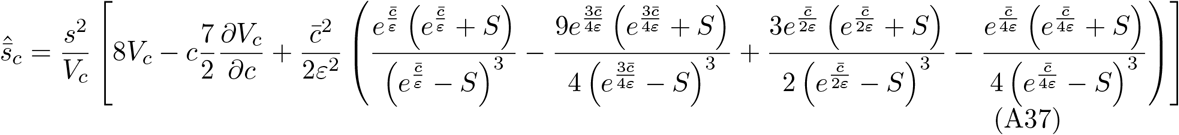

Making the variable change 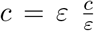 in Equations (A36) and (A37), so that it now depends of 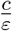 instead of *c*, we obtain that

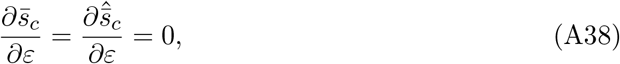

which demonstrates that 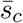 and 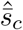 depend only on the ratio 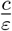. The same reasoning is followed for all three mutation types and will thus not be repeated.

###### Mutations altering *ε*

In this case, the leading order lifetime selection coefficient is

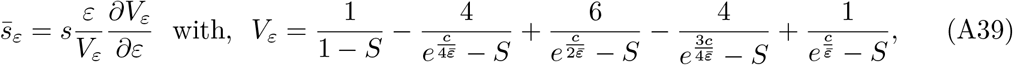

and the second order lifetime selection coefficient is given by

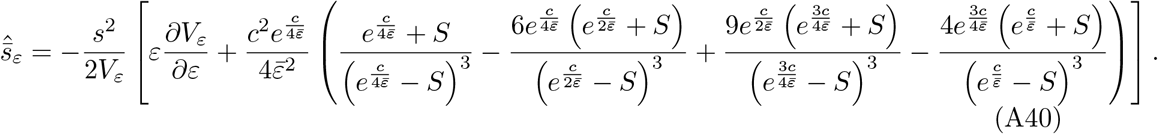

###### Mutations altering *S*

Finally, the leading order lifetime selection coefficient for mutations affecting survival is given by

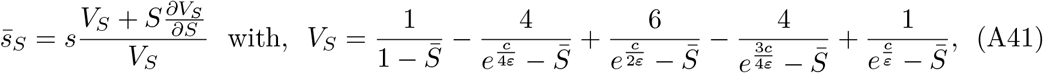

while the second order lifetime selection coefficient is

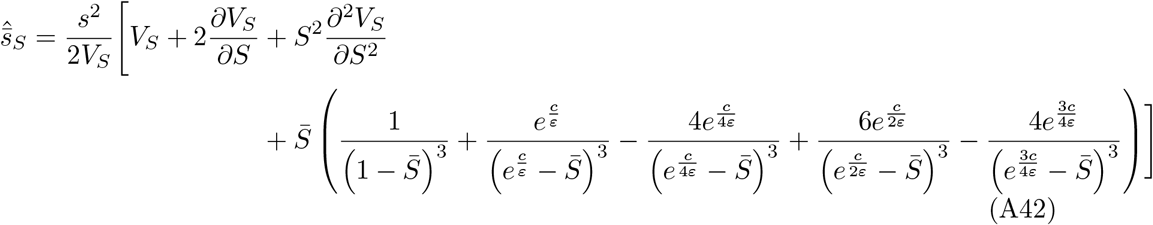

##### III.1.2 Quasi-linkage equilibrium results

To obtain an approximation for the number of mutations maintained at equilibrium, we need to find the point at which the amount of mutations entering the population due to mutation is compensated by the amount of mutations removed from the population by selection, that is the mutation-selection balance. To do so, one has to find the point at which the variation in allelic frequencies at the selected loci is worth zero. In the LF approach, selection is followed by reproduction, so that the change in allelic frequencies at the selected loci is given by Equations (A23) and (A28), that is

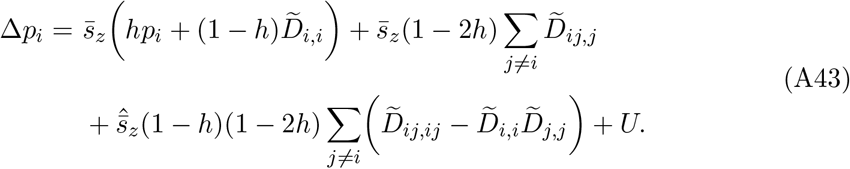

The leading order approximation of Equation (A43) is presented in Equation (8) in the main text. As mentioned in Appendix II.1, association 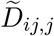 is of order *s*, so that the second term in Equation (A43) is of order *s*^2^, while the third term is of order 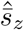, which is of order *s*^2^. Hence, both these terms can be neglected to leading order in *s* and Equation (A43) simplifies into

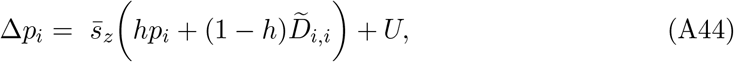

which corresponds to the first line of Equation (8) in the main text.

In order to solve solve Δ*p_i_* = 0, we need to obtain equilibrium expressions for genetic associations appearing in Equation (A43).

###### Equilibrium genetic associations under neutrality

As mentioned above, the effects of selection on 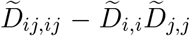 can be neglected, as it is generated by inbreeding even in the absence of selection. Assuming neutrality, neglecting terms in 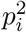, and using Equation (A29), we have

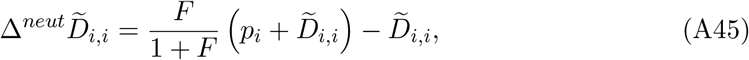

which at equilibrium yields

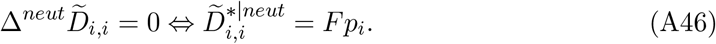

Similarly, using Equation (A31), we obtain

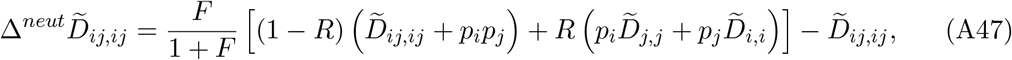

which assuming free recombination 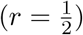 and using Equation (A46), yields

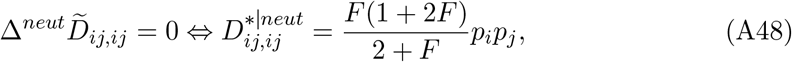

so that at equilibrium, the term 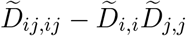 can be approximated by

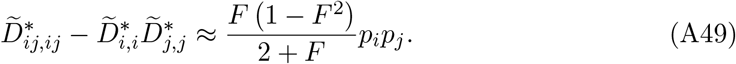

###### Equilibrium genetic associations to leading order in *s*

Using Equations (A26) and (A29), we have

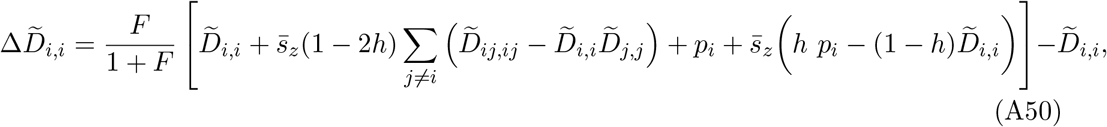

which, when replacing 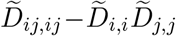 with Equation (A49), and neglecting direct selection effects (which is reasonable when the number of mutations segregating is large; Roze, 2015), yields

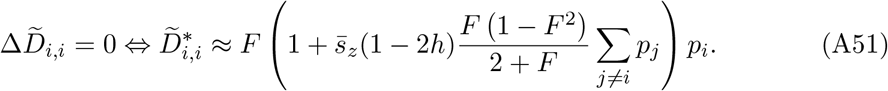

Finally, using Equations (A27) and (A30), and injecting Equation (A49), we have

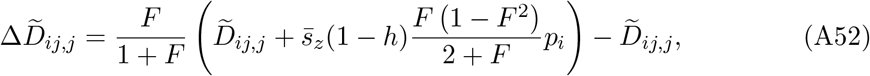

which yields, at equilibrium,

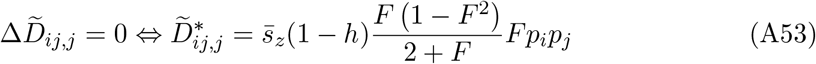

###### Mutation-selection balance

The number of mutations per haploid genome maintained at mutation selection balance is the solution of Δ*p_i_* = 0. However, even after injecting Equations (A49), (A51) and (A53) into Equation (A23), it still cannot be solved explicitly, as it also depends on *p_i_* through lifetime selection coefficients, which are functions of the mean trait of the population. Thus, we have to solve the system given by Equation (7) in the main text numerically, which we do in a *Wolfram Language Script*, that is the script version of a *Mathematica* notebook, available from GitHub (links are available from the journal office).

#### III.2 Life cycle approach

In the LC approach, selection at different stages is considered separately, thus we denote the selection coefficients associated with selection phase *k*, when mutations affect trait *z*, as 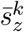 and 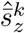 (Fig. S2). The stages at which selection occurs in our model are gametes production (denoted by *k* = *g*), juvenile recruitment (denoted *k* = *j*) and adult survival (*k* = *s*).

**Figure S2:**
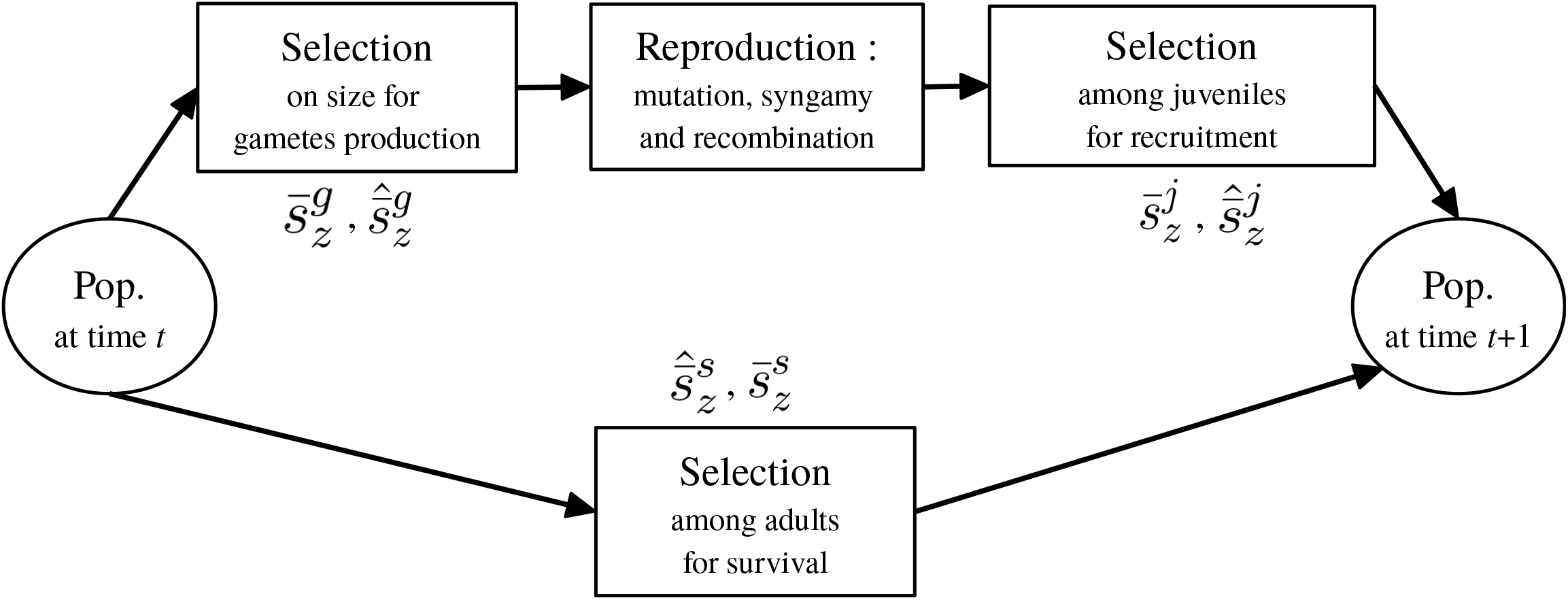
Steps followed over the course of one timestep with the LF approach

##### III.2.1 Stage specific selection coefficients

###### Gametes production

Selection on gametes production occurs in all three models (*k* = *g*) since fecundity is assumed to be proportional to size. Size at age *t* (*G_t_*) is the solution of Equation (2). It is given by

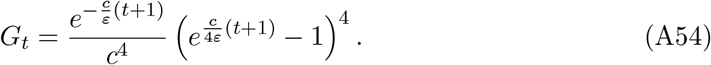

Thus, the average size of an individual given its genotype, which gives its fitness during this selection phase, is

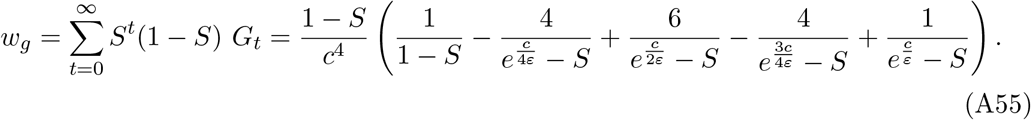

For any trait *z*, the selection coefficients at this stage, 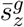 and 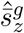 are obtained by applying Equation (A20) to Equation (A55).

###### Mutations affecting growth

In the case of mutations affecting growth, that is affecting the maintenance cost *c* or the production cost *ε*, these selection coefficients are equal to the lifetime selection coefficients presented in Appendix III.1, that is, they are given by Equations (A36) and (A37) for mutations affecting *c*, and Equations (A39) and (A40) for mutations affecting *ε*.

###### Mutations affecting survival

In the case of mutations affecting survival, the selection coefficients are given by

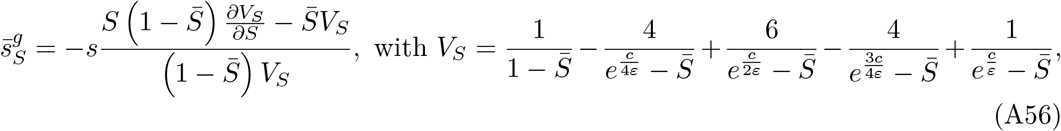

and,

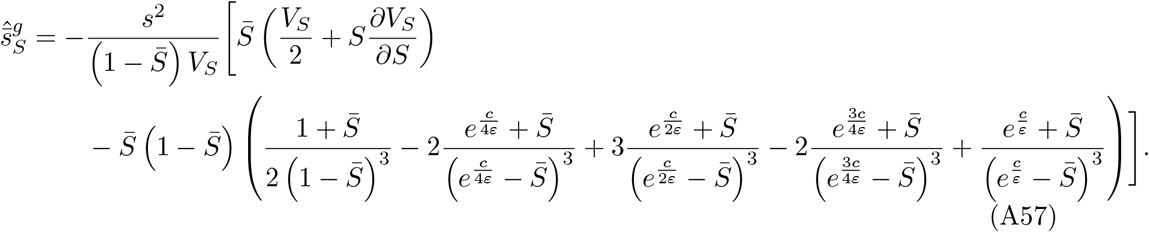

###### Survival and recruitment

Selection on survival occurs in juveniles (*k* = *j*) and in adults (*k* = *s*). Individual fitnesses during survival selection stages are given by their survival probability. Hence, when mutations affect the maintenance or the production cost (*i.e*. not survival), we have 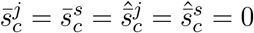 and 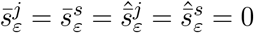. On the other hand, when mutations affect survival given by 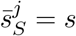 and 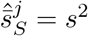 in juveniles, and 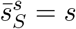 and 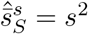 in adults.

##### III.2.2 Quasi-linkage equilibrium results

###### Allelic frequency change

In the LC approach, we distinguish between selection occuring in juveniles and in adults, so that the change in allelic frequency at the *i^th^* locus, Δ*p_i_*, is given by

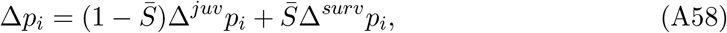

where 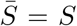 when mutations affect growth, and depends on the genetic composition of the population when they affect survival.

Among juveniles, we first have selection for gametes production, which is followed by reproduction, and selection for recruitment. Applying Equations (A23) and (A28), to these successive stages, one obtains, to second order in *s*,

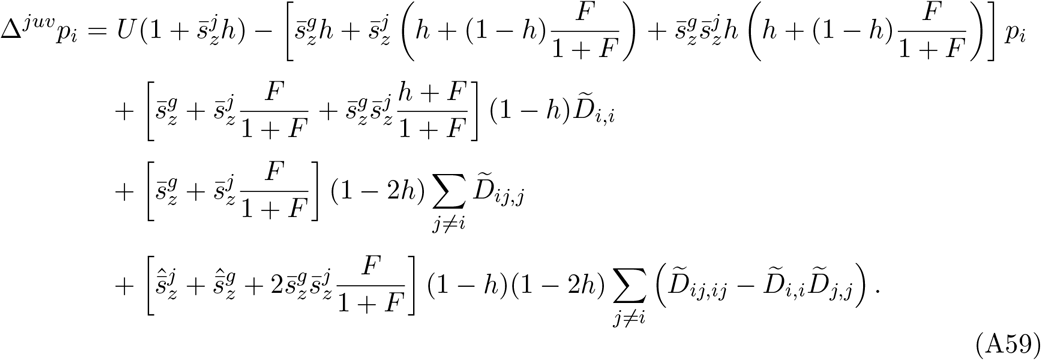

Besides, among surviving adults, only selection for survival occurs, so that to second order in *s* we have

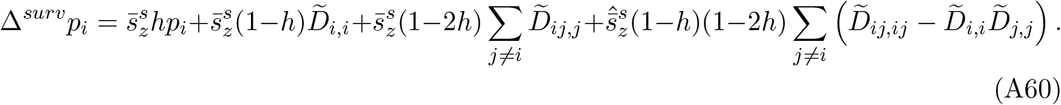

Putting Equations (A59) and (A60) together, this yields

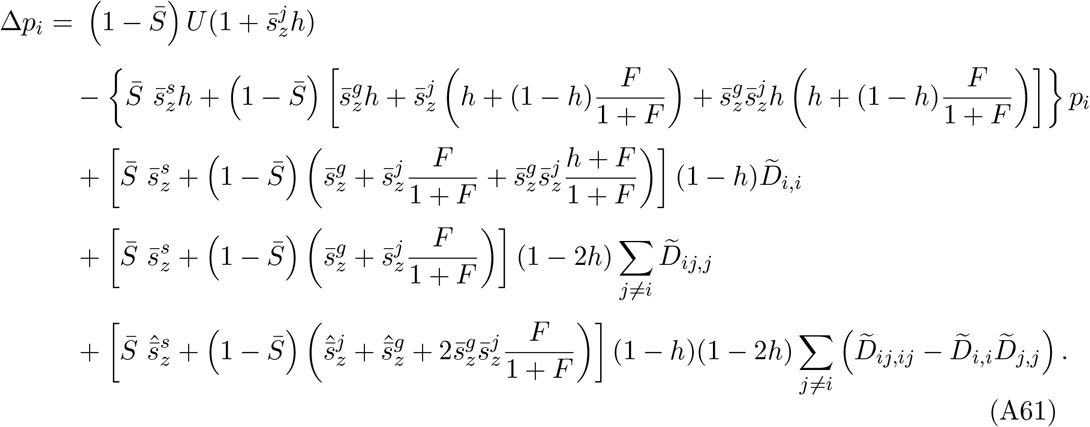

###### Genetic associations at quasi-linkage equilibrium

Similar to the LF approach, we know need to derive QLE approximations for genetic associations. In the case of associations derived under neutrality, the associations remain unchanged so that we still have

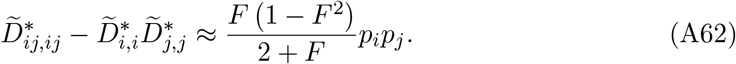

In the case of genetic associations affected by selection on the other hand, results differ from the LF approach. By the same logic as for allelic frequency change, the changes in these associations over the course of one timestep are given by

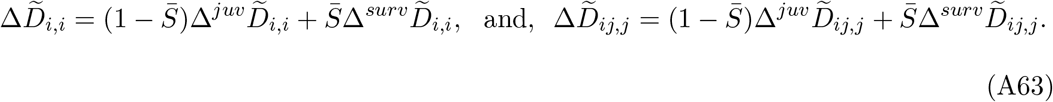

Applying the methods described in Appendix II to the case of association 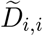, we have

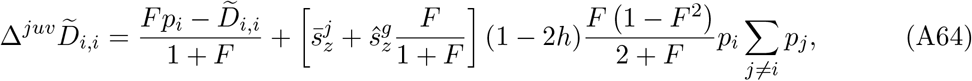

and,

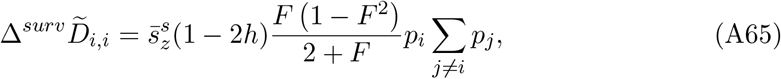

which yields

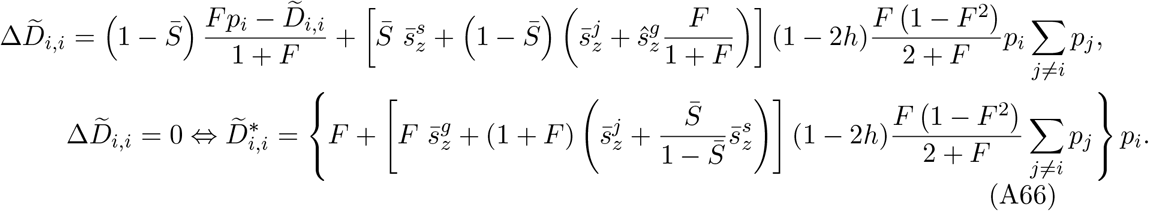

As for the 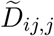 association, we have

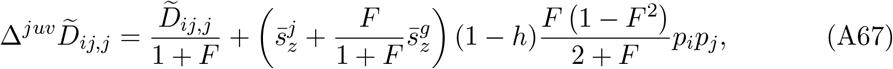

and,

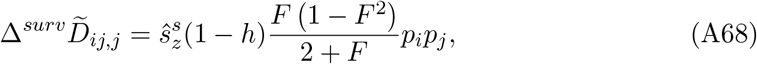

which yields

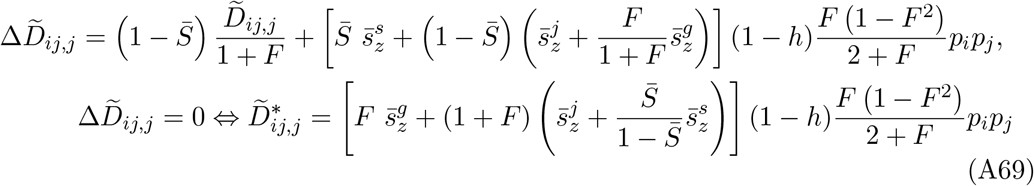

###### Mutation-selection balance

Solving Δ*p_i_* = 0 is done numerically in the same way as for the LF approach, using a *Wolfram Language Script* available from GitHub (links are available from the journal office).

#### III.3 Inbreeding depression and mutation load

In this section, we describe how an approximation for inbreeding depression and for the mutation load (Crow, 1958) can be obtained given the average number of mutations per haploid genome.

##### Inbreeding depression, *δ*

Inbreeding depression is defined as the relative decrease in lifetime fitness of selfed individuals relative to outcrossed ones (Charlesworth and Charlesworth, 1987). In our case, lifetime fitness is directly dependent individuals’ value for the trait *z*, through Equation (A35).

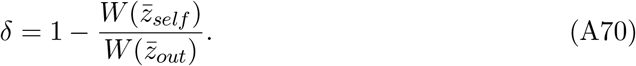

Thus, to compute an approximation for inbreeding depression, we need to obtain the average trait of selfed and outcrossed individuals, 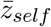 and 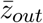. In the general population, the average of the trait is given by Equation (A19), that is

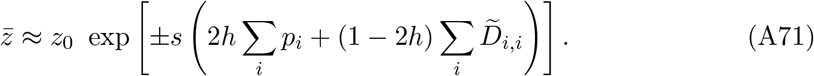

Among the outcrossed, we have 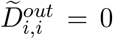 since they are produced by random mating. Hence,

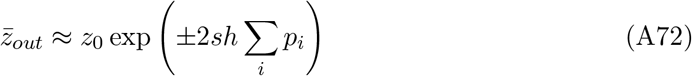

On the other hand, 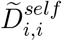 is given by

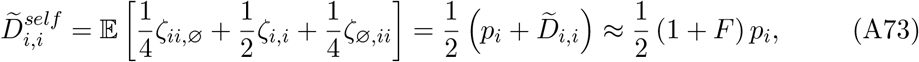

so that,

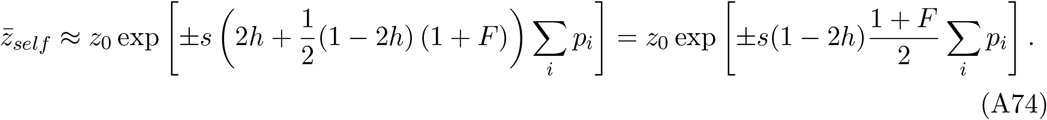

##### Mutation Load, *L*

The mutation load was defined by Crow (1958) as the decrease in mean fitness of the population compared to an optimal population bearing no mutation. Hence, in our model it is given by

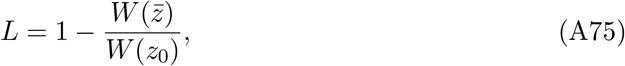

where *W*(*z*) is the lifetime fitness given by Equation (A35).

### IV Fitness landscapes

In Figure S3, lifetime fitness is presented as a function of phenotypic deviation Δ*P*. For each trait *z*, we have *z* = *z*_0_ (1 + Δ*P*). The resulting fitness landscape for each set of parameters is rescaled by its maximal value, which is always obtained for Δ*P* = 0.

**Figure S3:**
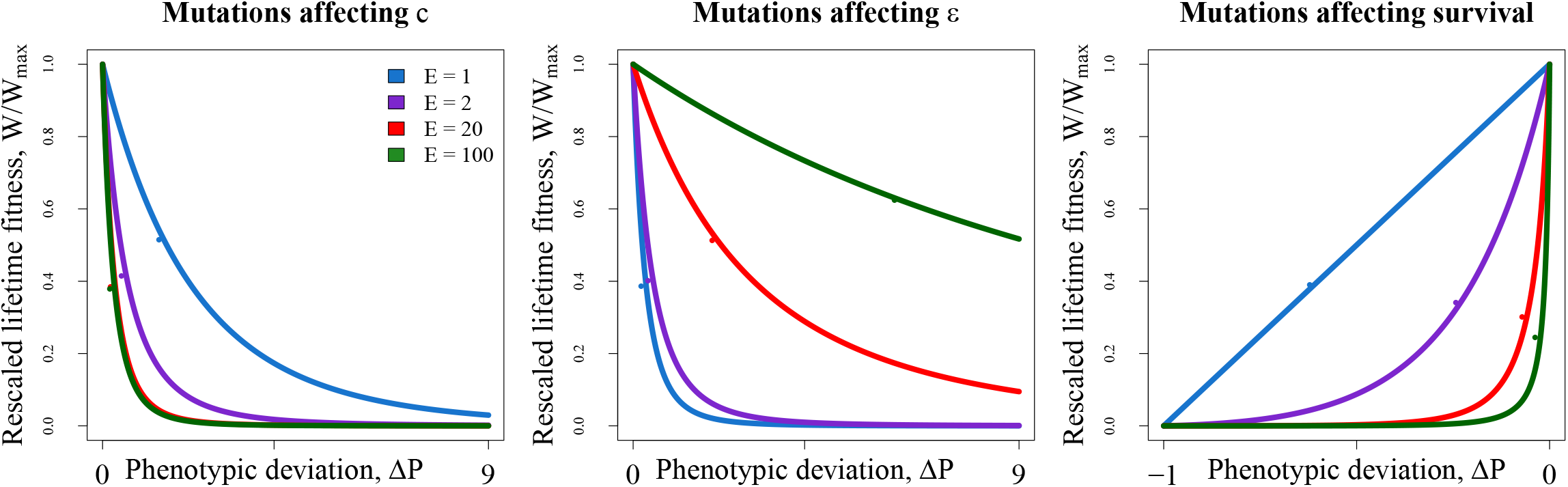
Rescaled fitness landscapes for each mutation type, for initial life expectancies (*E* = 1; 2; 20; 100). Lines depict fitness, while dots depict the mutation-selection equilibrium fitness reached for 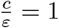, *h* = 0.25, *s* = 0.005, and *U* = 0.5.

### V Comparison of mean phenotypic deviation for different magnitude of effect of mutations

**Figure S4:**
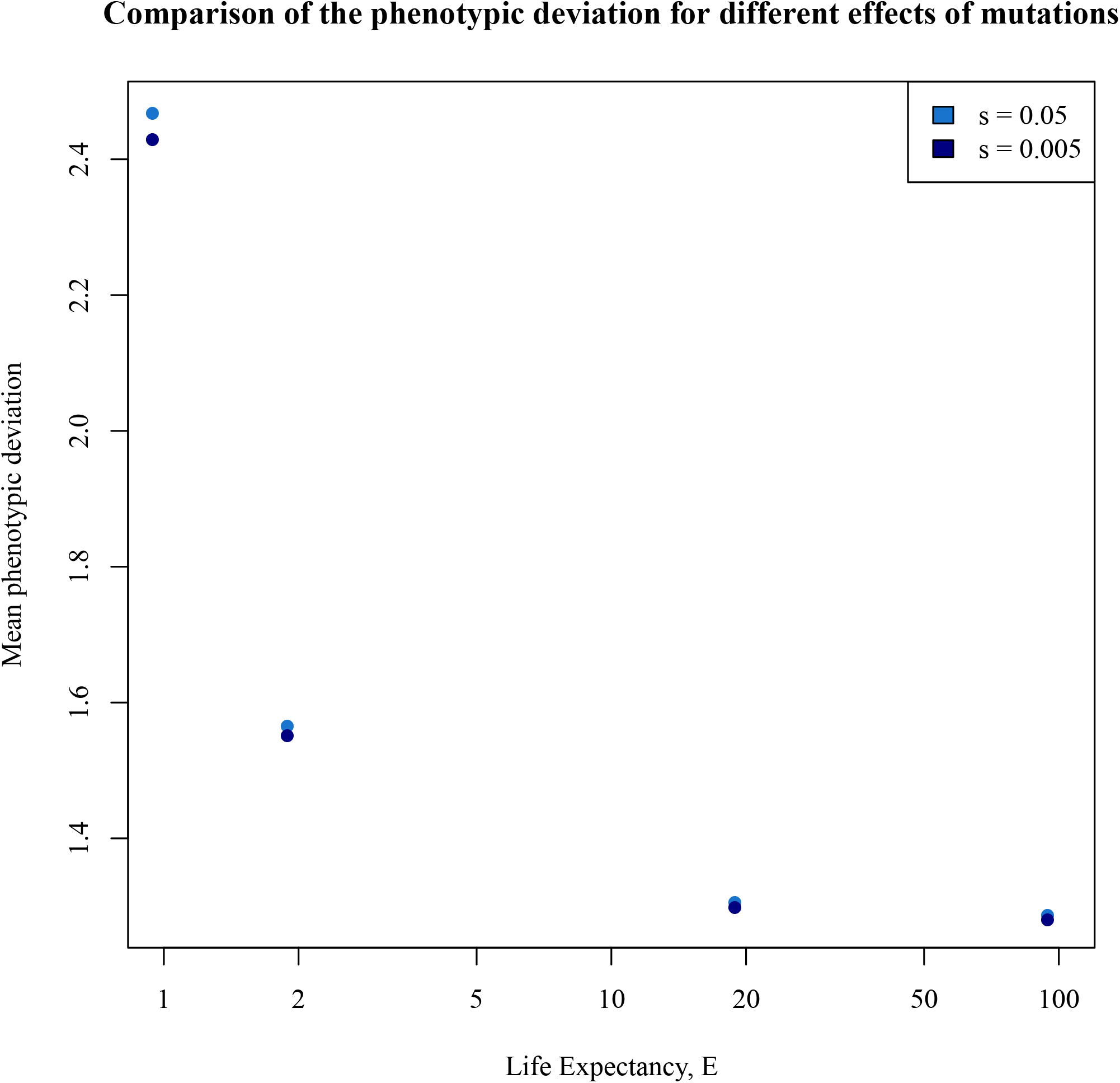
Comparison of the mean phenotypic deviation, for mutations affecting the maintenance cost (*c*), for two different magnitude of effect of mutations (*s* = 0.05 and *s* = 0.005). Parameters used are 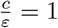, *h* = 0.25, *U* = 0.5.

### VI Mutations per haploid genome, inbreeding depression and mutation load for other parameter values

**Figure S5:**
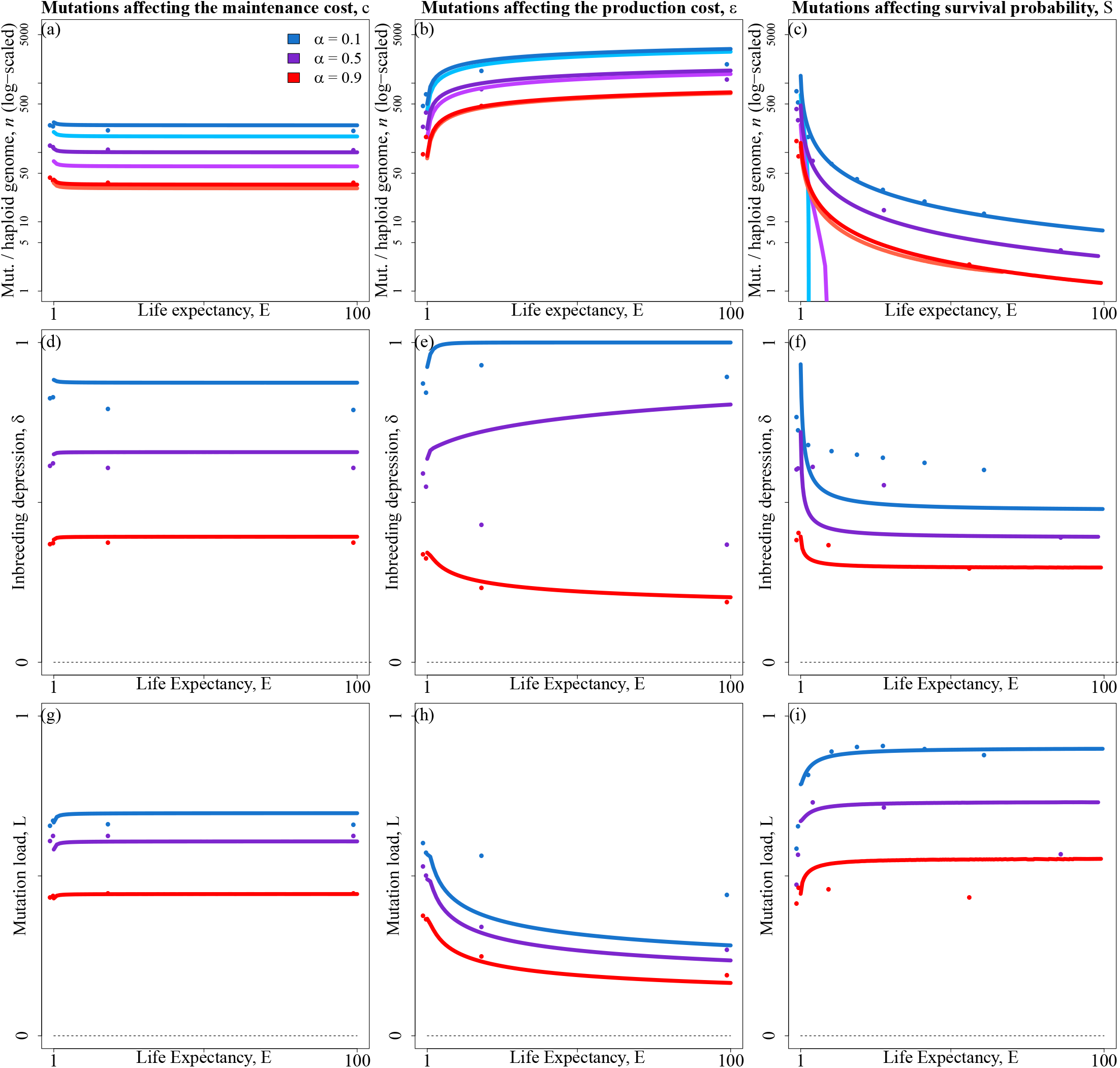
Average number of mutations per haploid genome (*n*, top row), inbreeding depression (*δ*, middle row), and mutation load (*L*, bottom row) as a function of life expectancy (*E*), for three selfing rates : *α* = 0.1 (blue), *α* = 0.5 (purple) and *α* = 0.9 (red). Each column corresponds to one type of mutation. Dots: simulation results. Dots: simulation results. Lighter lines: LF approach predictions. Darker lines: LC approach predictions. Parameters shown here are 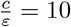, *U* = 0.5, *s* = 0.005, *h* = 0.10.

**Figure S6:**
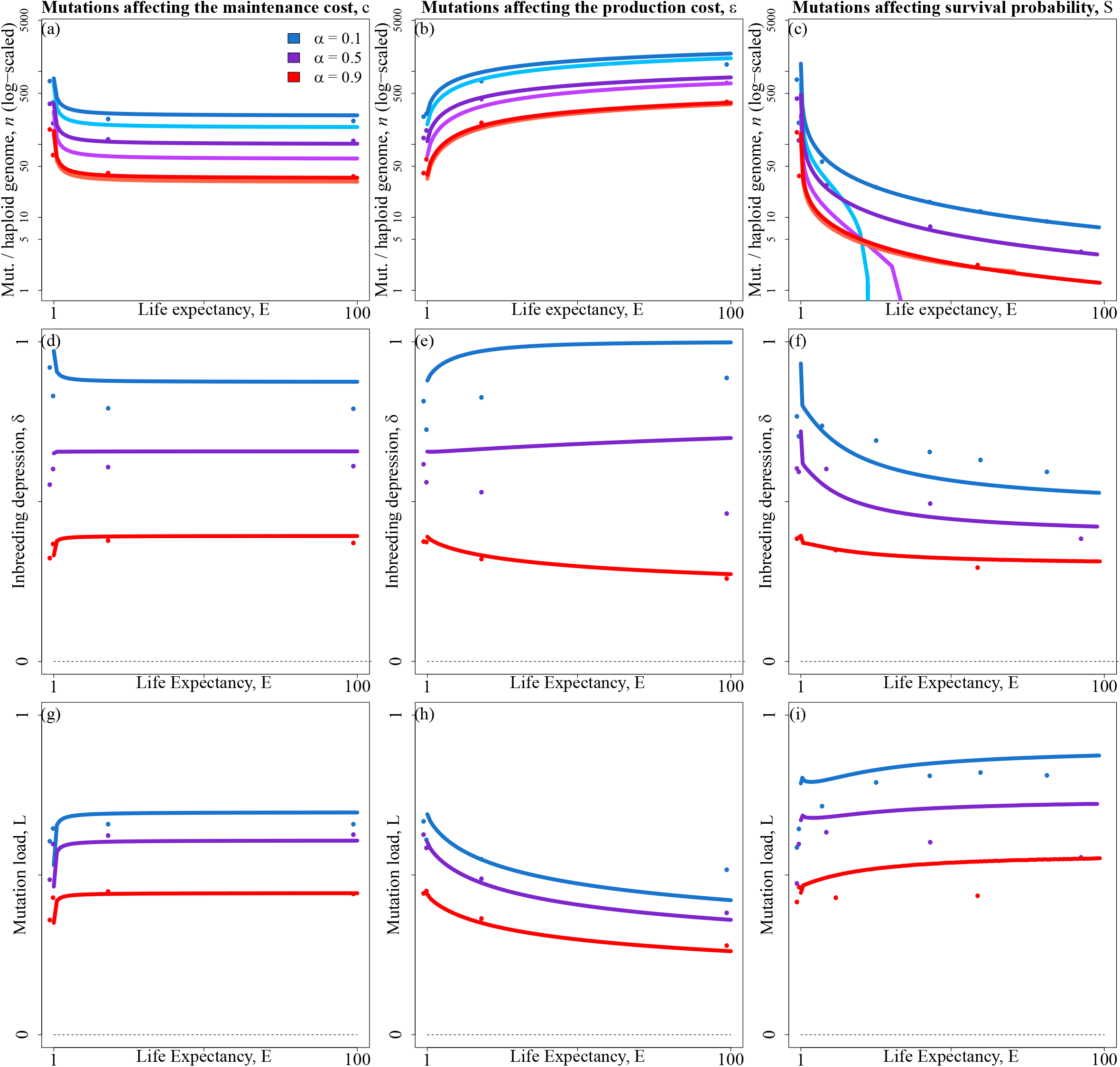
Average number of mutations per haploid genome (*n*, top row), inbreeding depression (*δ*, middle row), and mutation load (*L*, bottom row) as a function of life expectancy (*E*), for three selfing rates : *α* = 0.1 (blue), *α* = 0.5 (purple) and *α* = 0.9 (red). Each column corresponds to one type of mutation. Dots: simulation results. Dots: simulation results. Lighter lines: LF approach predictions. Darker lines: LC approach predictions. Parameters shown here are 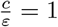, *U* = 0.5, *s* = 0.005, *h* = 0.10.

**Figure S7:**
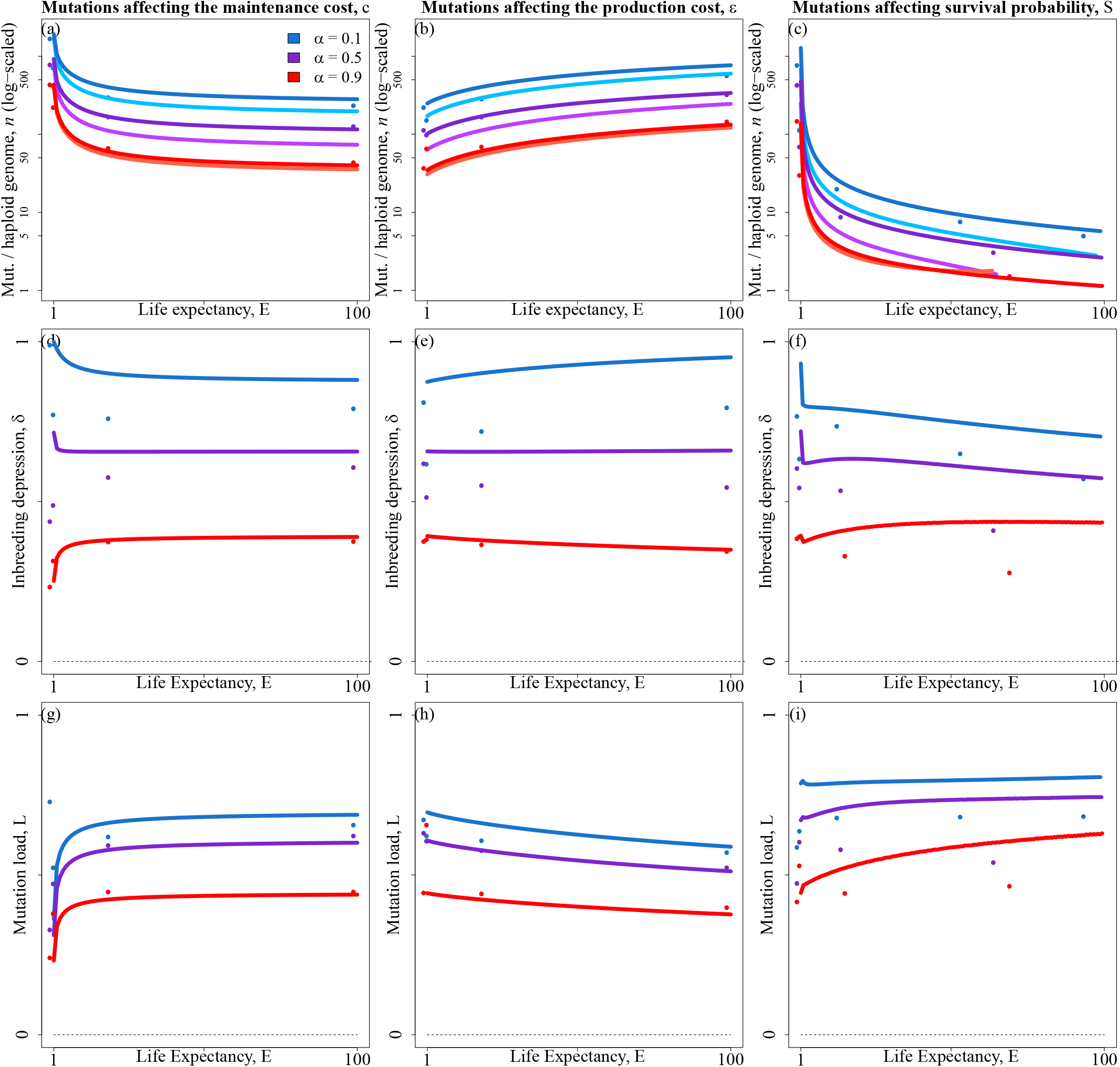
Average number of mutations per haploid genome (*n*, ntop row), inbreeding depression (*δ*, middle row), and mutation load (*L*, bottom row) as a function of life expectancy (*E*), for three selfing rates : *α* = 0.1 (blue), *α* = 0.5 (purple) and *α* = 0.9 (red). Each column corresponds to one type of mutation. Dots: simulation results. Dots: simulation results. Lighter lines: LF approach predictions. Darker lines: LC approach predictions. Parameters shown here are 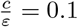, *U* = 0.5, *s* = 0.005, *h* = 0.10.

**Figure S8:**
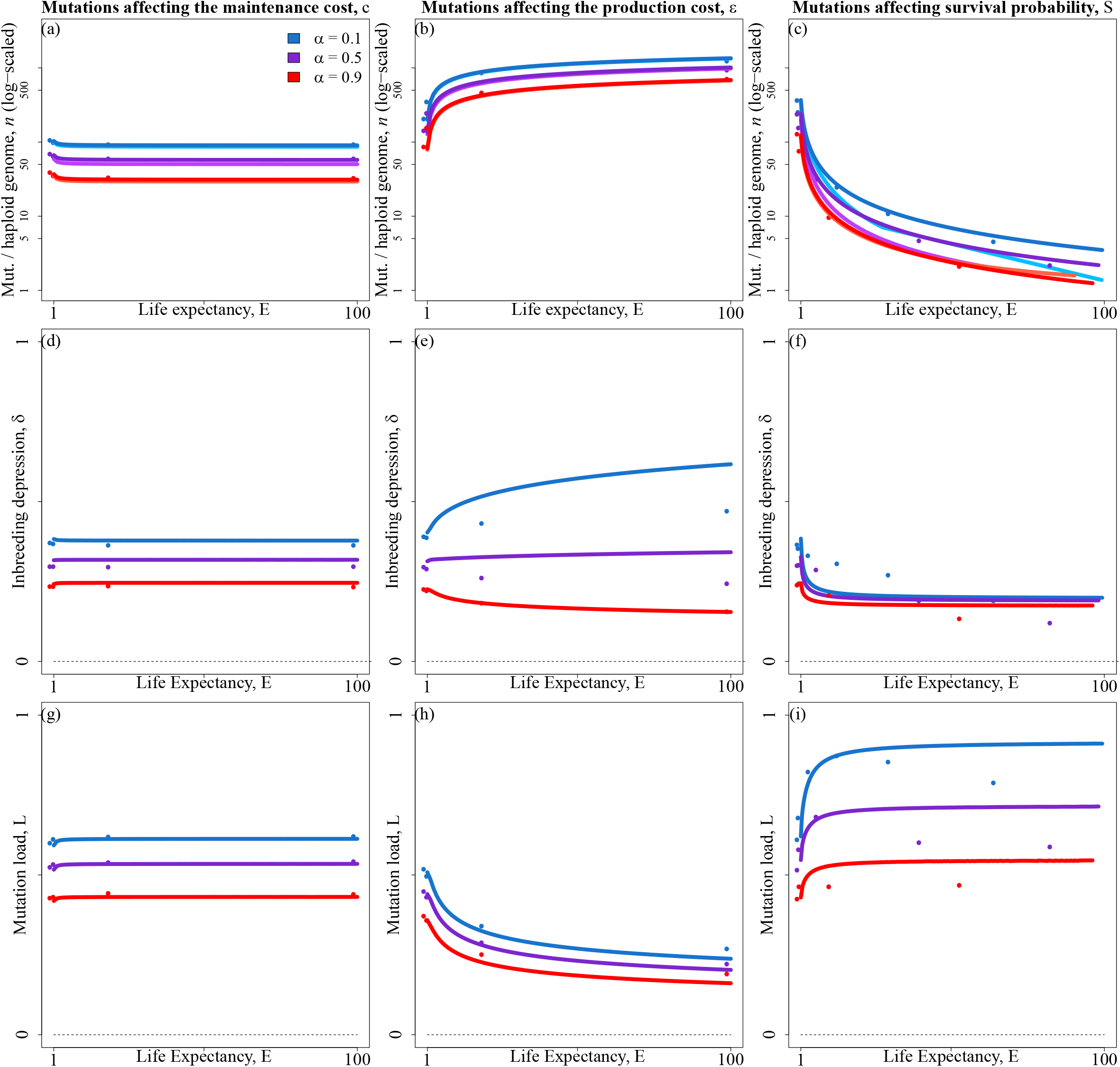
Average number of mutations per haploid genome (*n*, ntop row), inbreeding depression (*δ*, middle row), and mutation load (*L*, bottom row) as a function of life expectancy (*E*), for three selfing rates : *α* = 0.1 (blue), *α* = 0.5 (purple) and *α* = 0.9 (red). Each column corresponds to one type of mutation. Dots: simulation results. Dots: simulation results. Lighter lines: LF approach predictions. Darker lines: LC approach predictions. Parameters shown here are 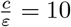, *U* = 0.5, *s* = 0.005, *h* = 0.25.

**Figure S9:**
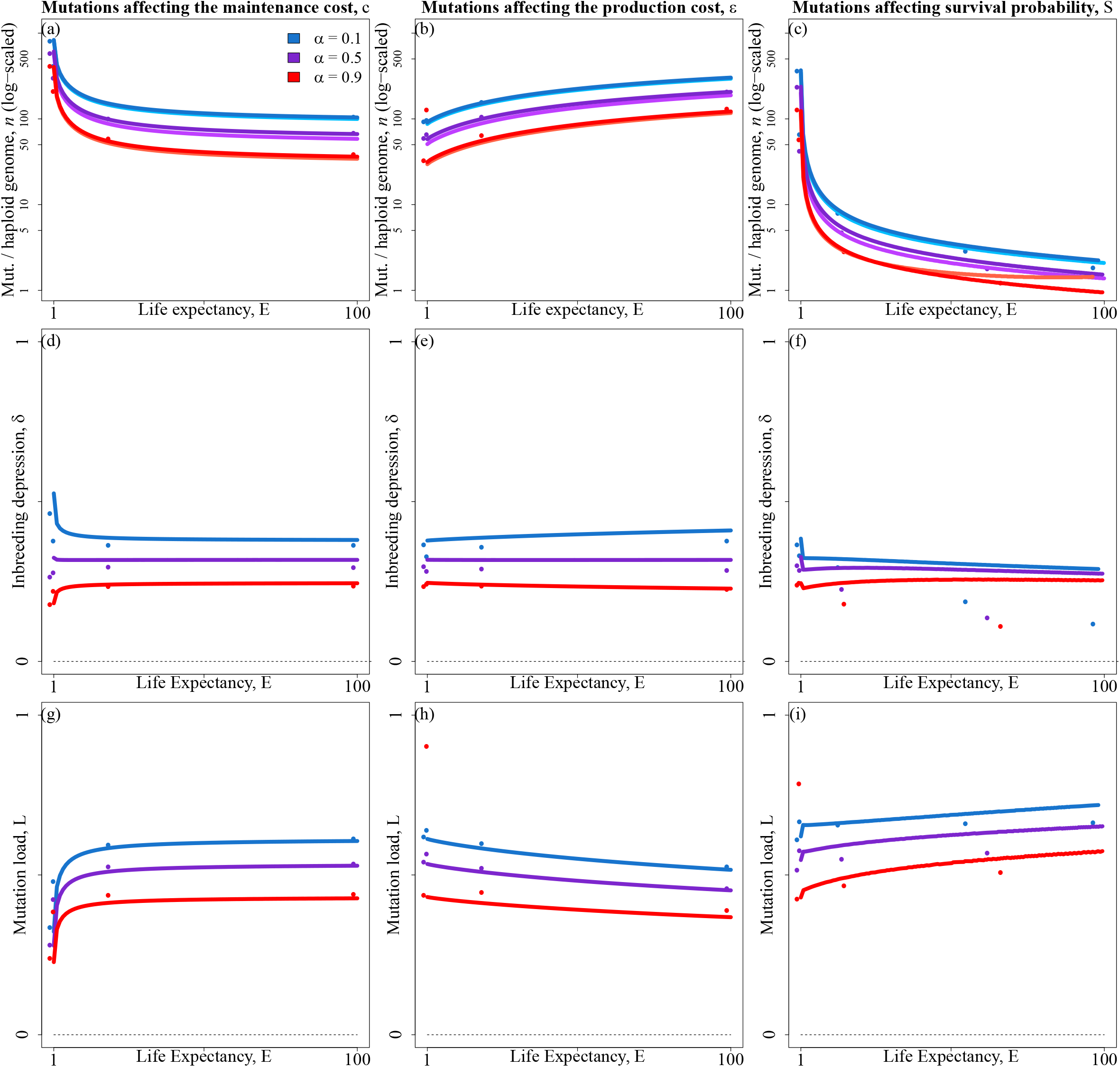
Average number of mutations per haploid genome (*n*, ntop row), inbreeding depression (*δ*, middle row), and mutation load (*L*, bottom row) as a function of life expectancy (*E*), for three selfing rates : *α* = 0.1 (blue), *α* = 0.5 (purple) and *α* = 0.9 (red). Each column corresponds to one type of mutation. Dots: simulation results. Dots: simulation results. Lighter lines: LF approach predictions. Darker lines: LC approach predictions. Parameters shown here are 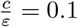, *U* = 0.5, *s* = 0.005, *h* = 0.250.

**Figure S10:**
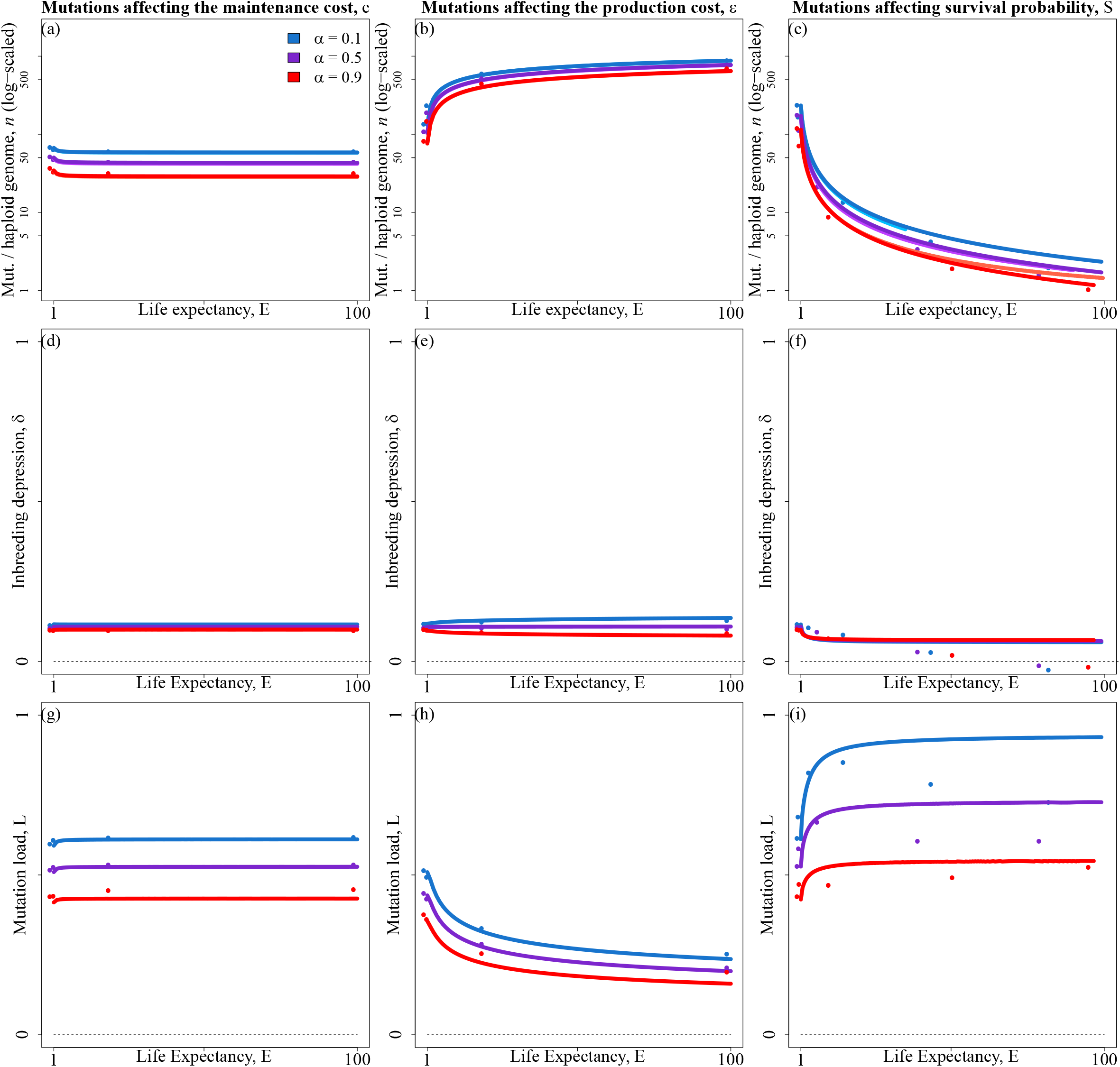
Average number of mutations per haploid genome (*n*, ntop row), inbreeding depression (*δ*, middle row), and mutation load (*L*, bottom row) as a function of life expectancy (*E*), for three selfing rates : *α* = 0.1 (blue), *α* = 0.5 (purple) and *α* = 0.9 (red). Each column corresponds to one type of mutation. Dots: simulation results. Dots: simulation results. Lighter lines: LF approach predictions. Darker lines: LC approach predictions. Parameters shown here are 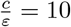, *U* = 0.5, *s* = 0.005, *h* = 0.40.

**Figure S11:**
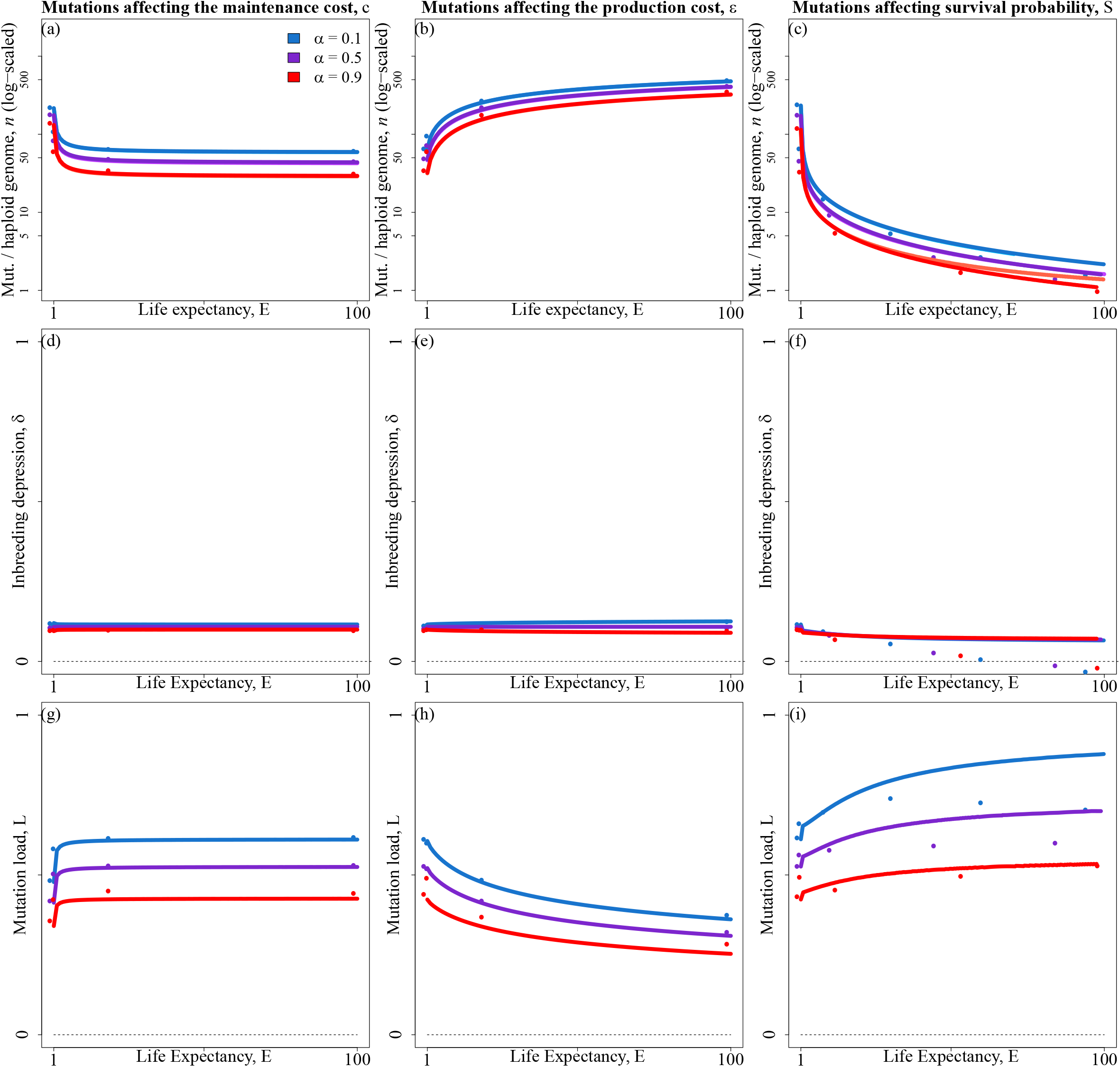
Average number of mutations per haploid genome (*n*, ntop row), inbreeding depression (*δ*, middle row), and mutation load (*L*, bottom row) as a function of life expectancy (*E*), for three selfing rates : *α* = 0.1 (blue), *α* = 0.5 (purple) and *α* = 0.9 (red). Each column corresponds to one type of mutation. Dots: simulation results. Dots: simulation results. Lighter lines: LF approach predictions. Darker lines: LC approach predictions. Parameters shown here are 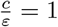, *U* = 0.5, *s* = 0.005, *h* = 0.40.

**Figure S12:**
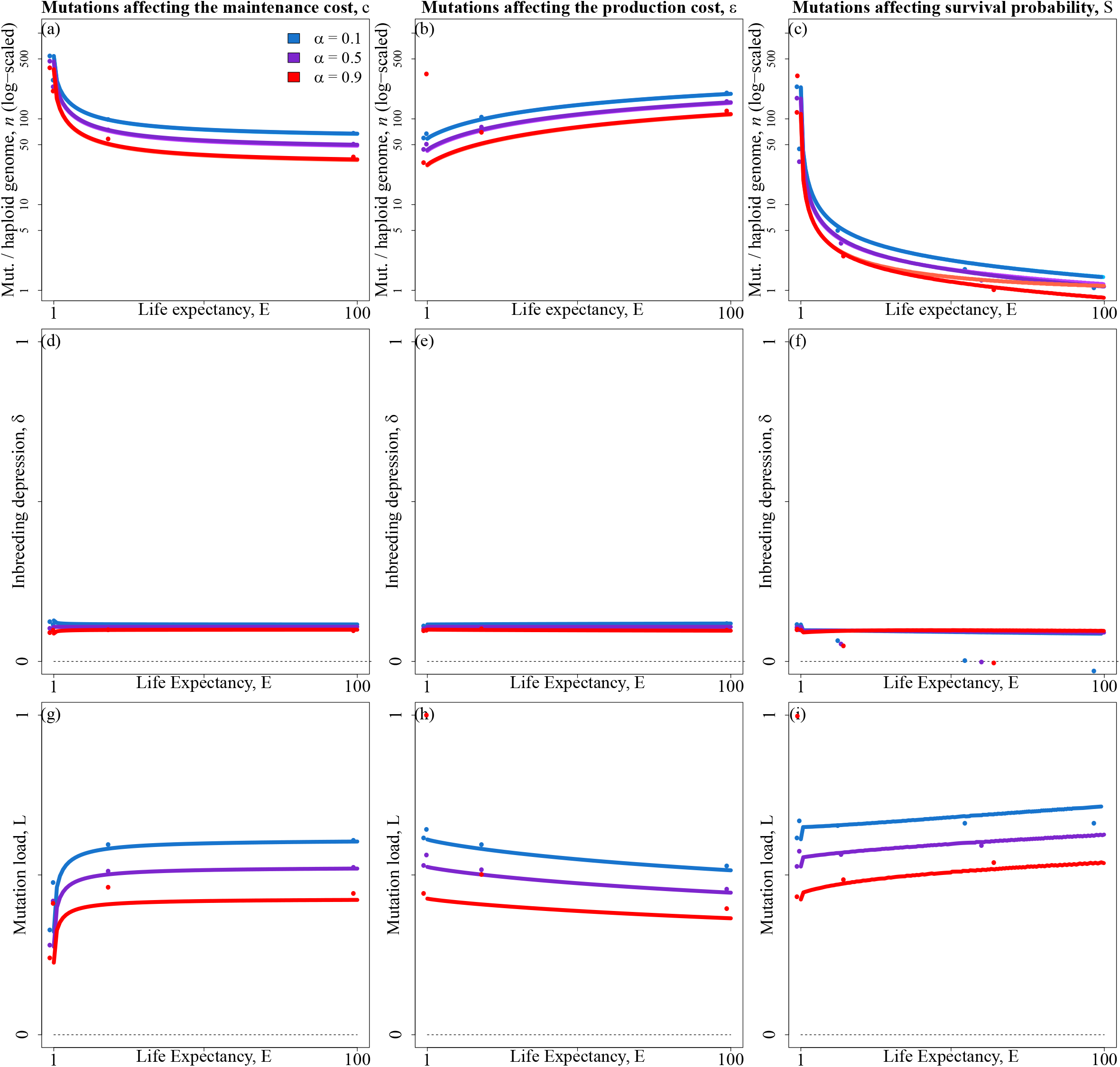
Average number of mutations per haploid genome (*n*, ntop row), inbreeding depression (*δ*, middle row), and mutation load (*L*, bottom row) as a function of life expectancy (*E*), for three selfing rates : *α* = 0.1 (blue), *α* = 0.5 (purple) and *α* = 0.9 (red). Each column corresponds to one type of mutation. Dots: simulation results. Dots: simulation results. Lighter lines: LF approach predictions. Darker lines: LC approach predictions. Parameters shown here are 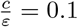, *U* = 0.5, *s* = 0.005, *h* = 0.40.

## References

Abu Awad, D. and Roze, D. (2018). Effects of partial selfing on the equilibrium genetic variance, mutation load and inbreeding depression under stabilizing selection. Evolution, 72:751–769.

Abu Awad, D. and Roze, D. (2019). Epistasis, inbreeding depression and the evolution of self-fertilization. BioRXiv.

Angeloni, F., Ouborg, N., and Leimu, R. (2011). Meta-analysis on the association of population size and life history with inbreeding depression in plants. Biological Conservation, 144:35–43.

Barrett, S. C. and Harder, L. D. (1996). The comparative biology of pollination and mating in flowering plants. Philosophical Transactions of the Royal Society of London B, 351:1271–1280.

Barton, N. and Turelli, M. (1991). Natural and sexual selection on many loci. Genetics, 127:229–255.

Bataillon, T. and Kirkpatrick, M. (2000). Inbreeding depression due to mildly deleterious mutations in finite populations: size does matter. Genetics Research, 75:75–81.

Bobiwash, K., Schultz, S., and Schoen, D. (2013). Somatic deleterious mutation rate in a woody plant: estimation from phenotypic data. Heredity, 111:338–344.

Burian, A., Barbier de Reuille, P., and Kuhlemeier, C. (2016). Patterns of stem cell divisions contribute to plant longevity. Current Biology, 26:1385–1394.

Charlesworth, B. (1980). Evolution in age-structured populations. Cambridge Studies in Mathematical Biology.

Charlesworth, B., Morgan, M., and Charlesworth, D. (1991). Multilocus models of inbreeding depression with synergistic selection and partial self-fertilization. Genetics, 57:177–194.

Charlesworth, D. and Charlesworth, B. (1987). Inbreeding depression and its evolutionary consequences. Annual Review of Ecology and Systematics, 18:237–268.

Charlesworth, D., Morgan, M., and Charlesworth, B. (1990). Inbreeding depression, genetic load, and the evolution of outcrossing rates in a multilocus system with no linkage. Evolution, 44:1469–1489.

Charlesworth, D. and Willis, J. (2009). The genetics of inbreeding depression. Nature reviews genetics, 10:783–796.

Crow, J. (1958). Some possibilities for measuring selection intensities in man. Human Biology, 61:763–775.

DeHaan, L. and Van Tassel, D. (2014). Useful insights from evolutionary biology for developing perennial grain crops. American Journal of Botany, 101:1801–1819.

Duminil, J., Hardy, O., and Petit, R. (2009). Plant traits correlated with generation time directly affect inbreeding depression and mating system and indirectly genetic structure. BMC Evolutionary Biology, 9:177.

Duputié, A. and Massol, F. (2013). An empiricist’s guide to theoretical predictions on the evolution of dispersal. Interface Focus, 40:20130028.

Ehrlén, J. and Lehtilä, K. (2002). How perennial are perennial plants? Oikos, 98:1070–1072.

Enquist, B., Brown, J., and West, G. (1998). Allometric scaling of plant energetics and population density. Nature, 395:163–165.

Epinat, G. and Lenormand, T. (2009). The evolution of assortative mating and selfing with in- and outbreeding depression. Evolution, 63:2047–2060.

Franco, M. and Silvertown, J. (1996). Life-history variation in plants: an exploration of the fast-slow continuum hypothesis. Philosophical Transactions of the Royal Society of London B, 351:1341–1348.

Gros, P.-A., Le Nagard, H., and Tenaillon, O. (2009). The evolution of epistasis and its links with genetic robustness, complexity and drift in a phenotypic model of adaptation. Genetics, 182:277–293.

Husband, B. and Schemske, D. W. (1996). Evolution of the magnitude and timing of inbreeding depression in plants. Evolution, 50:54–70.

Kirkpatrick, M., Johnson, T., and Barton, N. (2002). General models of multilocus evolution. Genetics, 161:1727–1750.

Klinkhamer, P., Meelis, E., De Jong, T., and Weiner, J. (1985). On the analysis of sizedependent reproductive output in plants. Functional Ecology, 6:308–316.

Kondrashov, A. (1985). Deleterious mutations as an evolutionary factor. II. facultative apomixis and selfing. Genetics, 111:635–653.

Lanfear, R. (2018). Do plants have a segregated germline? PloS Biology, 16.

Morgan, M. (2001). Consequences of life history for inbreeding depression and mating system evolution in plants. Proceedings of the Royal Society of London B, 268:1817–1824.

Munoz, F., Violle, C., and Cheptou, P.-O. (2016). CSR ecological strategies and plant mating systems: outcrossing increases with competitiveness but stress-tolerance is related to mixed mating. Oikos, 125:1296–1303.

Otto, S. and Orive, M. (1995). Evolutionary consequences of mutation and selection within an individual. Genetics, 141:1173–1187.

Peters, R. (1983). The ecological implications of body size. Cambridge studies in ecology.

Petit, R. and Hampe, A. (2006). Some evolutionary consequences of being a tree. Annual Review of Ecology, Evolution, and Systematics, 37:187–214.

Phillips, P. (2008). Epistasis—the essential role of gene interactions in the structure and evolution of genetic systems. Nature Reviews Genetics, pages 855–867.

Plomion, C., Aury, J.-M., Amselem, J., Leroy, T., Murat, F., Duplessis, S., Faye, S., Francillonne, N., Labadie, K., Le Provost, G., Lesur, I., Bartholomé, J., Faivre-Rampant, P., Kohler, A., Leplé, J.-C., Chantret, N., Chen, J., Diévrat, A., Alaeitabar, T., Barbe, V., Belser, C., Bergès, H., Bodénès, C., Bogeat-Triboulot, M.-B., Bouffaud, M.-L., Brachi, B., Chancerel, E., Cohen, D., Couloux, A., Da Silva, C., Dossat, C., Ehrenmann, F. Gaspin, C., Grima-Pettenati, J., Guichoux, E., Hecker, A., Herrmann, S., Hugueney, P., Hummel, I., Klopp, C., Lalanne, C., Lascoux, M., Lasserre, E., Lemainque, A. Desprez-Loustau, M.-L., Luyten, I., Madoui, M.-A., Mangenot, S., Marchal, C., Mau-mus, F., Mercier, J., Michotey, C., Panaud, O., Picault, N., Rouhier, N., Rué, O., Rustenholz, C., Salin, F., Soler, M., Tarkka, M., Velt, A., Zanne, A., Martin, F., Wincker, P., Quesneville, H., Kremer, A., and Salse, J. (2018). Oak genome reveals facets of long lifespan. Nature Plants, 4:440–452.

Roze, D. (2015). Effects of interference between selected loci on the mutation load, inbreeding depression, and heterosis. Genetics, 201:745–757.

Roze, D. and Blanckaert, A. (2014). Epistasis, pleiotropy, and the mutation load in sexual and asexual populations. Evolution, 86:137–149.

Roze, D. and Michod, R. (2010). Deleterious mutations and selection for sex in finite, diploid populations. Genetics, 184:1095–1112.

Roze, D. and Rousset, F. (2005). Inbreeding depression and the evolution of dispersal rates: a multilocus model. The Amrican Naturalist, 166:708–721.

Salguero-Gómez, R., Jones, O. R., Jongejans, E., Blomberg, S. P., Hodgson, D. J., Mbeau-Ache, C., Zuidema, P. A., de Kroon, H., and Buckley, Y. M. (2016). Fast–slow continuum and reproductive strategies structure plant life-history variation worldwide. Proceedings of the National Academy of Sciences, 113(1):230–235.

Schmid-Siegert, E., Sarkar, N., Iseli, C., Calderon, S., Gouhier-Darimont, C., Chrast, J., Cattaneo, P., Schütz, F., Farinelli, L., Pagni, M., Schneider, M., Voumard, J., Jaboyed-off, L., Fankhauser, C., Hardtke, C., Keller, L., Pannell, J., Reymond, A., Robinson-Rechavi, M., Xenarios, I., and Reymond, P. (2017). Low number of fixed somatic mutations in a long-lived oak tree. Nature Plants, 12:926–929.

Schoen, D. and Schultz, S. (2019). Somatic mutation and evolution in plants. Annual Review of Ecology, Evolution and Systematics, 50:2.1–2.25.

Scofield, D. and Schultz, S. (2006). Mitosis, stature and evolution of plant mating systems: low-*ϕ* and high-*ϕ* plants. Proceedings of the Royal Society of London B, 273:275–282.

Stearns, S. (1992). The Evolution of Life Histories. Oxford University Press.

Watson, J., Platzer, A., Kazda, A., Akimcheva, S., Valuchova, S., Nizhynska, V., Nordborg, M., and Riha, K. (2016). Germline replications and somatic mutation accumulation are independent of vegetative life span in arabidopsis. Proceedings of the National Academy of Sciences, 113:12226–12231.

Weiner, J., Campbell, L., Pino, J., and Echarte, L. (2009). The allometry of reproduction within plant populations. Journal of Ecology, 97:1220–1233.

West, G., Brown, J., and Enquist, B. (1997). A general model for the origin of allometric scaling laws in biology. Science, 276:122–126.

West, G., Brown, J., and Enquist, B. (1999). A general model for the structure, function and allometry of plant vascular systems. Nature, 400:664–667.

West, G., Brown, J., and Enquist, B. (2001). A general model for ontogenetic growth. Nature, 413:628–631.

Winn, A., Elle, E., Kalisz, S., Cheptou, P.-O., Eckert, C., Goodwillie, C., Johnston, M., Moeller, D., Sargent, R., and Vallejo-Marìn, M. (2011). Analysis of inbreeding depression in mixed-mating plants provides evidence for selective interference and stable mixed mating. Evolution, 65:3339–3359.

